# Hyperosmotic stress induces rewiring of 3D chromatin interactions

**DOI:** 10.64898/2025.12.19.695003

**Authors:** J.P. Flores, Andrea A. Perreault, Zachary A. Drum, Chenxi Xu, Doris Cruz Z, Tooba Rashid, Jeremy D. Burgess, Geoffrey C. Fox, Gelila Petros, Yijia Wu, Ivana Y. Quiroga-Barber, HyunAh Kim, Isha Sahasrabudhe, Ye Jin Lee, Ethan Black, Yihuan Li, Justin Demmerle, Brian D. Strahl, Jill M. Dowen, Gang Greg Wang, Danfeng Cai, Douglas H. Phanstiel

## Abstract

Cells continually face environmental stressors that challenge homeostasis, yet how three-dimensional (3D) chromatin structure contributes to these stress responses remains unclear under hyperosmotic conditions. To investigate this, we map 3D chromatin structure, its molecular drivers, and transcriptional outcomes during sorbitol-induced hyperosmotic stress. Within 1 hour of sorbitol treatment, pre-existing loops and domains collapse, while several hundred de novo sorbitol-induced loops emerge. These loops are punctate, longer-range, and transient. 3D chromatin structure largely returns to baseline conditions by 24 hours. Loop reorganization is consistent across cell types and hyperosmotic stimuli. Sorbitol-induced loops require cohesin but not CTCF, and anchors are enriched at active promoters harboring GC-rich transcription factor motifs. Genes at these anchors show delayed activation following loop formation, consistent with loops acting upstream of transcription. Together, hyperosmotic stress triggers rapid, reversible, CTCF-independent chromatin reorganization that correlates with transcriptional adaptation.

---

Cells are constantly challenged by fluctuations in their environment, and their ability to mount rapid, coordinated responses to these stressors is critical for maintaining homeostasis and survival**^1–5^**. One common stressor encountered by human cells is rapid changes in osmolarity that must be adjusted to in order to preserve protein stability, membrane integrity, and nuclear organization**^6–10,10–12^**. In the kidney, for example, cells in the renal medulla experience high NaCl and urea and rely on osmoprotective programs to prevent damage, while in the immune system, elevated NaCl in the tumor microenvironment can enhance CD8^+^ T cell metabolic fitness and cytotoxicity**^10,13,14^**. Thus, hyperosmotic stress shapes cellular physiology in diverse tissues**^9,10^** and understanding how cells adapt to osmotic stress could therefore provide broadly relevant insights into human biology and the mechanisms of gene regulation.

While much of the response to hyperosmotic stress has been defined through its transcriptional outputs**^15–19^**, the upstream layers of regulation—particularly those involving three-dimensional (3D) chromatin structure—remain incompletely understood. However, 3D chromatin structure plays an important role in gene regulation, and mounting evidence suggests that the genome is rapidly remodeled when cells encounter hyperosmotic stress. We found that YAP1/TAZ, TEAD1, and accessible chromatin regions colocalize in human kidney cells under sorbitol-induced hyperosmotic stress, which suggests large-scale changes to 3D chromatin architecture**^20^**. Hyperosmotic stress has also been linked to redistribution of transcriptional co-regulators to accessible chromatin sites, weakening of chromatin domain boundaries, disruption of long-range looping interactions, widespread RNA polymerase II run-off and readthrough transcription, as well as large-scale chromatin condensation and compartment shifts**^18,19,21–25^**. In some systems, chromatin reorganization appears to be directed by specific architectural proteins rather than passive compaction alone**^8,26^**. Together, these studies suggest that hyperosmotic stress triggers coordinated structural and transcriptional remodeling of the genome.

Despite these findings, the connection between 3D chromatin structure and hyperosmotic stress response is not entirely clear. Many existing studies either did not directly map chromatin interactions, were performed in non-mammalian systems, or relied on shallow sequencing that could miss nuanced changes in 3D chromatin architecture**^19–21^**. As a result, the scope of 3D chromatin changes during hyperosmotic stress, the molecular machinery responsible, and the functional consequences for gene regulation remain incompletely defined.

To address these gaps, we mapped 3D chromatin structure, protein-DNA interactions, and gene expression in human cells responding to sorbitol-induced hyperosmotic stress. Together, we find rapid but transient remodeling of chromatin structure during hyperosmotic stress response and identify molecular features that may drive these structural changes. Our findings provide insights into how cells reorganize their nuclear architecture under stress and inform our understanding of the interplay between 3D chromatin structure and transcription in health and disease.

## Results

### Hyperosmotic stress triggers widespread rewiring of chromatin loops

To determine how hyperosmotic stress impacts 3D chromatin structure, we performed deeply sequenced Hi-C (8.18 billion total sequencing reads across 4 replicates per condition) in human embryonic kidney (HEK293T eGFP-YAP1) cells treated with 200 mM sorbitol for 1 hour, alongside untreated controls. Cells expressing eGFP-tagged YAP1 were used in order to follow up our recent studies mapping YAP1 phase separation in response to sorbitol, but as shown later, this modification did not influence results**^20^**. Hi-C contact maps revealed extensive structural changes after sorbitol treatment exemplified at loci shown in **Figure 1A**. The vast majority of pre-existing chromatin loops and chromatin structures were lost and were replaced via the formation of a smaller number of sorbitol-induced loops. Differential analysis revealed 14,714 lost and 361 sorbitol-induced loops after 1 hour of treatment (DESeq2, padj < 0.1) (**Figure 1B**). The global loss of pre-existing 3D chromatin structures has been observed in cells treated with 110mM NaCl**^19^**, but to the best of our knowledge, the gained loops in response to hyperosmotic stress had not been previously described. The vast majority (97.2%) of sorbitol-induced loops were formed *de novo* and were not observed in untreated cells, indicating a major reorganization of 3D contacts rather than strengthening of pre-existing loops **(Figure S1)**. Notably, however, approximately half (49%) of sorbitol-induced loop anchors overlap with anchor positions present in untreated cells **(Figure S1)**, suggesting that sorbitol-induced loops frequently repurpose existing architectural nodes.

**Fig. 1.**
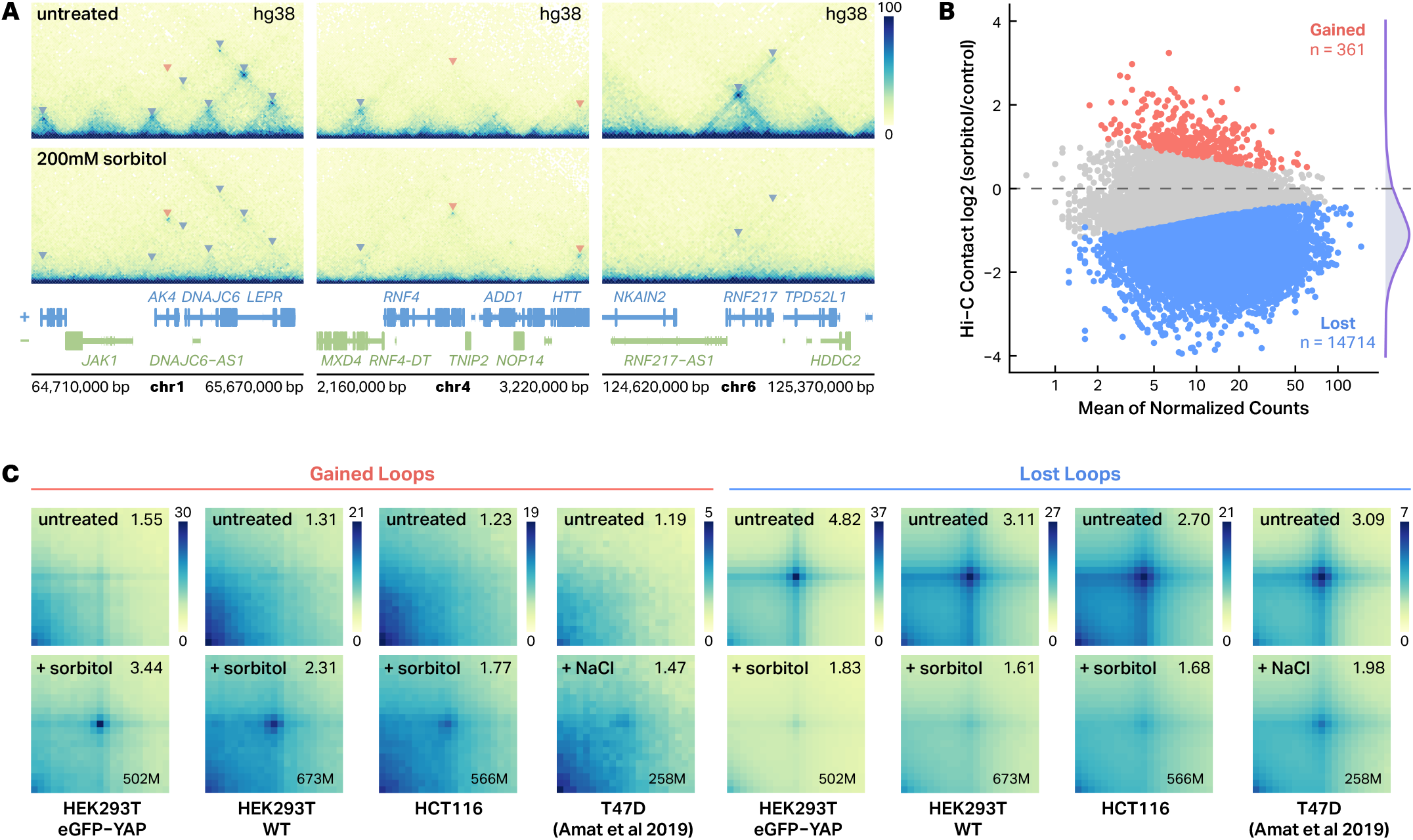
Hyperosmotic stress induces large-scale rewiring of chromatin interactions. **A**, Representative Hi-C contact maps showing chromatin structure changes at selected genomic loci (hg38) in HEK293T eGFP-YAP1 cells. Top panels show untreated controls, bottom panels show cells treated with 200 mM sorbitol for 1 hour. Triangular indicators mark chromatin loops (red triangles = gained loops, blue triangles = lost loops). **B**, MA plot showing differential chromatin loop analysis between sorbitol-treated and untreated HEK293 cells. Red points indicate significantly gained loops (n = 361), blue points indicate significantly lost loops (n = 14,714). Statistical significance was determined by DESeq2 analysis (padj < 0.1, |log**2**FC| > 0 for gained loops, |log**2**FC| < 0 for lost loops). Density plot (purple) on the right show the distribution log2 fold changes. **C**, Aggregate Peak Analysis (APA) heatmaps demonstrating loop formation and loss across multiple cell lines and stimuli. Each heatmap shows aggregate contact frequency centered on loop anchors, with APA scores (foreground/background enrichment) indicated in the upper-right corner of each matrix. Contact frequency is normalized by the max value for each cell type across treatment conditions. Treatment conditions: sorbitol (200 mM, 1h) for HEK293T eGFP-YAP1, HEK293T WT, and HCT116; NaCl (110 mM) for T47D. Average sequencing read depths (in millions) are indicated in the lower-right corner of each APA matrix.

To assess the generality of these effects, we performed Hi-C in both wild-type (WT) HEK293T cells and the colorectal cancer line, HCT116. To evaluate changes in looping across datasets, we performed aggregate peak analysis (APA) using the same set of loop anchor coordinates defined from the HEK293T eGFP-YAP1 differential loop calls as a shared reference. This approach enabled a direct, cell type-independent comparison of loop signal at a consistent set of genomic positions. In all three cell lines, APA showed strong depletion of pre-existing loop signal and the emergence of new loops following sorbitol treatment (**Figure 1C**). To determine if these results were due to sorbitol specifically or to hyperosmotic stress in general, we reanalyzed Hi-C data describing the treatment of the breast cancer cell line T47D with 110mM NaCl. While the initial paper describing those results did not report any gained loops, aggregate peak analysis revealed evidence of sorbitol-induced looping at the same regions where we observed gained loops in HEK293T WT, HEK293T eGFP-YAP1, and HCT116 cells. Weaker enrichment was observed in cells treated with NaCl than sorbitol, which could indicate differences in loop strength, loop locations, or the shallower sequencing of the NaCl-treated data-set (0.51 billion reads). Regardless, the persistence of loop enrichment across all conditions highlights the robustness of hyperosmotic stress-induced chromatin rewiring (**Figure 1C**). These findings demonstrate that hyperosmotic stress induces widespread rewiring of chromatin loops independent of cell type or specific osmoregulating stimuli.

Because hyperosmotic stress can alter nuclear architecture at multiple length scales, we next asked whether sorbitol-induced chromatin rewiring was accompanied by broader changes in genome organization and nuclear morphology. Hi-C analysis revealed weakened chromatin domain insulation and reduced genome-wide A/B compartment strength in both eGFP-YAP1 and YAPΔTAD cells, indicating broad disruption of higher-order chromatin organization that occurs independently of YAP1 phase separation (Figure S2). We then quantified nuclear volume from three-dimensional confocal image stacks before and after sorbitol treatment. Sorbitol-treated cells had significantly smaller nuclei than untreated controls (Fig. S3). Together, these findings suggest that hyperosmotic stress induces global nuclear compaction while simultaneously disrupting canonical features of 3D genome organization.

### Sorbitol-induced loops are transient and characteristically distinct from pre-existing loops

To gain insight into the mechanisms driving sorbitol-induced loop formation, we characterized the contact frequency profiles of sorbitol-induced loops. To do so, we standardized loop dimensions to a uniform 150kb width and built aggregate Hi-C maps for 241 medium to long-range (>= 150kb) sorbitol-induced loops at 10kb resolution **(Figure 2A, top)**. For comparison, we built aggregate Hi-C maps for a matched set of pre-existing loops (**Figure 2A, middle**), selected to have similar loop sizes and contact frequencies as sorbitol-induced loops. Sorbitol-induced loops were more punctate than pre-existing loops (**Figure 2B, bottom left**), even when controlling for loop strength and distance (**Figure S5)**. The contact domains within sorbitol-induced loops also exhibited reduced contact frequency and had weaker ‘stripes’ along their borders. Sorbitol-induced loops are also longer on average than pre-existing loops **(Figure S1C)**. The exact implications of these differences are unclear but are consistent with differences in the mechanisms of formation or maintenance compared to the pre-existing loops.

**Fig. 2.**
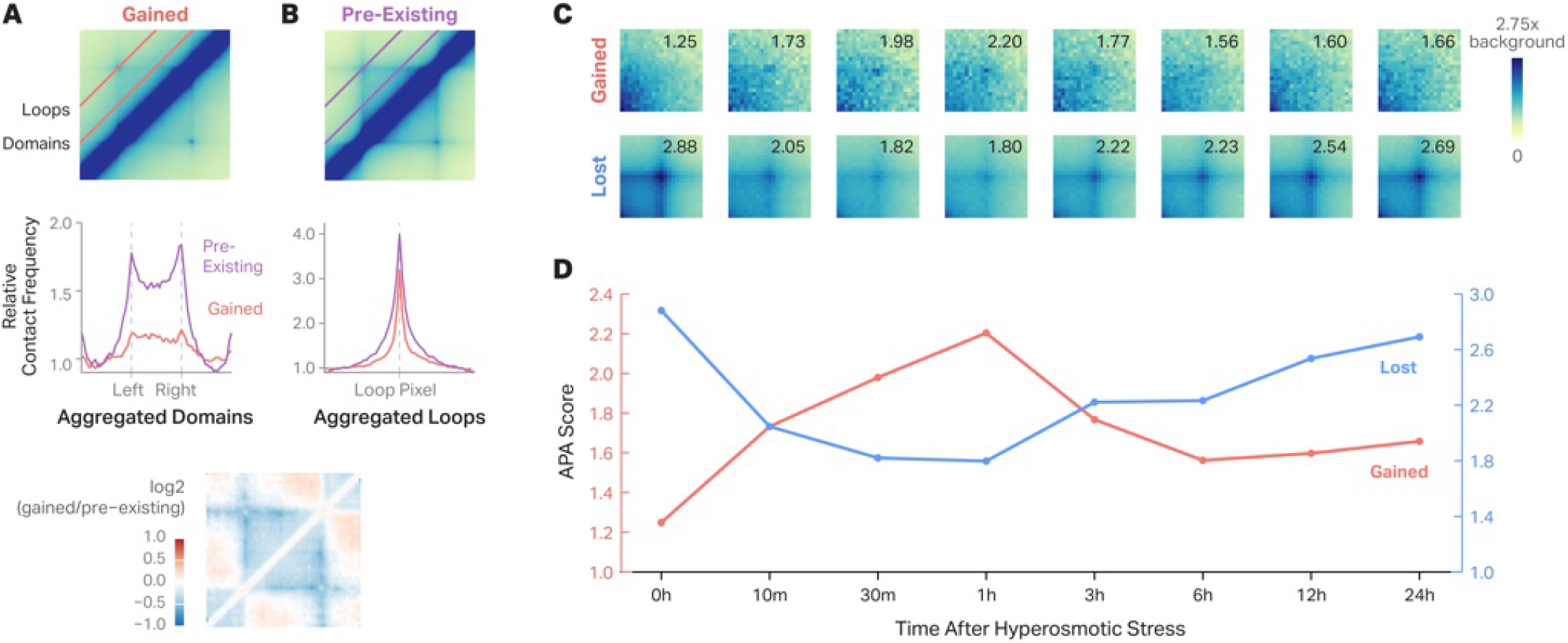
Sorbitol-induced loops are more punctate, form weaker chromatin domains, and peak at 1 hour of treatment. **A**, Aggregate Hi-C contact maps comparing gained and pre-existing loops and domains, standardized to a uniform 150-kb width (top). The red and purple lines on the plot denote the pixels represented in the line plots below. Line plots show the relative contact frequency within the aggregated domain boundaries of gained (red line) and pre-existing domains (purple line) (middle). A differential log**2**(gained/pre-existing) contact map is shown, highlighting the reduced domain insulation surrounding gained loops relative to pre-existing loops (bottom). **B**, Aggregate Hi-C contact maps comparing gained (top) and pre-existing loops (bottom). Line plots below display the relative contact frequency profiles across aggregated chromatin loops before and after treatment. **C**, Aggregate Hi-C time-course analysis of chromatin loops following hyperosmotic stress. Heatmaps show aggregate contact patterns for gained loops (top row) and lost loops (bottom row) at eight time points: 0h, 10m, 30m, 1h, 3h, 6h, 12h, and 24h after sorbitol treatment. APA scores are indicated in the upper-right corner of each matrix. Color scale represents background-normalized contact frequency (0 to 2.75×), where background is defined as the median contact value from the top-left and bottom-right corners of each aggregate matrix. **D**, Quantitative analysis of chromatin loop enrichment over time. APA scores (y-axis) plotted against time after hyperosmotic stress treatment (x-axis: 0h to 24h). Red line shows gained loops, blue line shows lost loops.

To understand the timing of formation and the persistence of sorbitol-induced loops, we quantified the dynamics of sorbitol-induced looping across an 8-point Hi-C (0h control, 10min, 30min, 1h, 3h, 6h, 12h, and 24h) time course. The cells were maintained in 200 mM sorbitol for the entire time course. Aggregate Peak Analysis (APA) revealed that sorbitol-induced loops are detectable by the earliest measured time point (10 min) after treatment and are most prominent at 1 hour. Sorbitol-induced loops are weakened but retain higher contact frequency than baseline even at 24 hours. Preexisting loops were disrupted within 10 minutes of sorbitol treatment but began to re-establish as early as 3 hours after treatment **(Figure 2C–D)**. Together, these results suggest that sorbitol triggers a distinct class of chromatin loops that are punctate, transient, and form weaker chromatin domains.

### CTCF and cohesin are retained at sorbitol-induced loop anchors

To determine whether stress-induced chromatin rewiring involves changes in architectural protein occupancy, we performed CTCF and RAD21 CUT&Tag in eGFP-YAP1 HEK293T cells that were either untreated or treated with sorbitol for 1 hour. Consistent with previous findings**^19^**, genome-wide differential analysis revealed that hyperosmotic stress led to global decreases in binding intensity (**Figure 3A**) with 13,742 CTCF peaks and 204 RAD21 peaks exhibiting reduced occupancy and only 421 CTCF peaks and 42 RAD21 peaks exhibiting increased occupancy (**Figure S6,** DESeq2, padj < 0.05 for CTCF and YAP1; padj < 0.1 for RAD21). Consistent with these changes in chromatin occupancy, fluorescence imaging revealed a marked redistribution of CTCF-EGFP and RAD21-mEmerald within the nucleus following sorbitol treatment (**Figure S7**). However, decreases in CTCF occupancy differed across gained and lost loops. Loops that were lost in response to sorbitol exhibited a precipitous loss in both CTCF and RAD21 occupancy which explains their decreased contact frequency (**Figure 3B**). In contrast, over 70% of gained loop anchors were bound by CTCF even before sorbitol treatment and largely remained bound after sorbitol treatment (**Figure 3B**). Similar trends were observed for RAD21 occupancy. Moreover, CTCF and RAD21 binding sites that were between gained loop anchors exhibited a more pronounced decrease in occupancy in response to sorbitol than binding sites at gained loop anchors (**Figure 3C & S4**). This selective retention of CTCF binding and RAD21 occupancy at sorbitol-induced loop anchors could explain the changes observed in chromatin looping in response to sorbitol. These results are exemplified in **Figure 3D** in which lost CTCF and RAD21 binding is associated with lost loops and retained CTCF and RAD21 binding is observed at gained loop anchors.

**Fig. 3.**
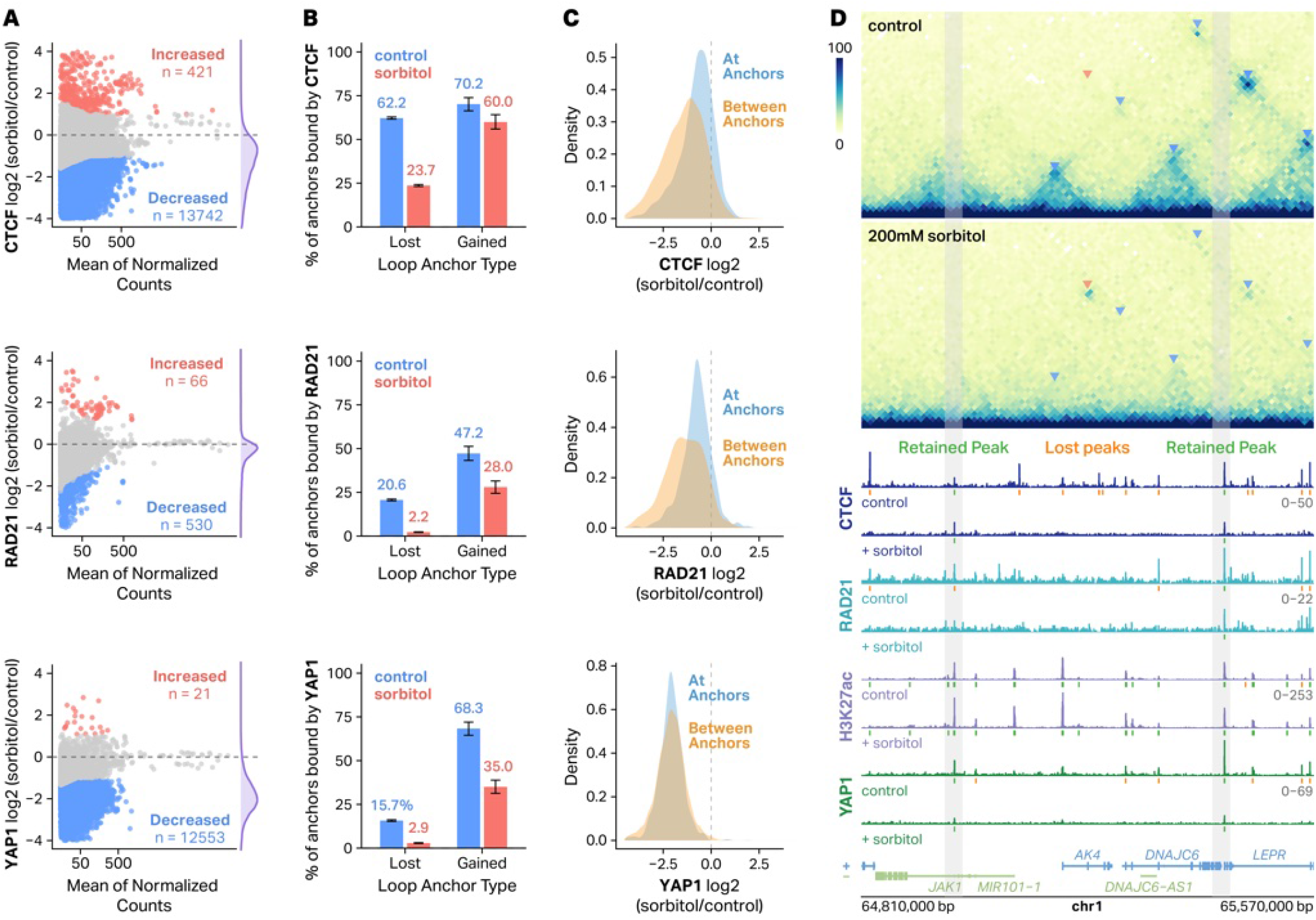
CTCF, cohesin, and YAP1 are retained at gained loop anchors following hyperosmotic stress. **A**, MA plots showing differential analysis of CTCF (top), RAD21 (middle), and YAP1 (bottom) binding (log2FC sorbitol/control) versus mean of normalized counts. Sites with significantly increased or decreased (DESeq2, padj < 0.05 for CTCF and YAP1; padj < 0.1 for RAD21) binding are highlighted in red and blue, respectively. **B**, Barplots depicting the percentage of loop anchors bound by CTCF (top), RAD21 (middle), and YAP1 (bottom). **C**, Density plots showing the distribution of changes in CTCF (top), RAD21 (middle), and YAP1 (bottom) occupancy of peaks at loop anchors (blue) compared to those within loop anchors (orange). **D**, Hi-C contact maps (10-kb resolution, hg38) at an example locus on chromosome 1 showing loop formation after 200 mM sorbitol treatment. Arrows denote gained (red) or lost (blue) loops. Corresponding CTCF, RAD21, H3K27ac, and YAP1 CUT&Tag tracks are shown below. Retained (green) and lost (orange) peaks are denoted.

We next asked what sequence and chromatin features could potentially explain the differences we observed in CTCF retention in response to sorbitol treatment. Analysis of CUT&Tag signal as a function of baseline occupancy revealed that higher-occupancy CTCF and RAD21 sites exhibited more moderate decreases in occupancy following sorbitol treatment compared to lower occupancy sites **(Figure S8A)**. Retained CTCF peaks had significantly higher CTCF motif position weight matrix (PWM) scores than lost CTCF peaks **(Figure S8B)**. RAD21 was significantly less depleted at retained CTCF sites compared to lost CTCF peaks **(Figure S8C)**, suggesting that these retained CTCF sites might function as loop extrusion boundaries. Retained CTCF peaks were also more likely to overlap gene promoters **(Figure S8D)**. Taken together, these results suggest that high occupancy CTCF peaks, with strong CTCF motifs, at or near gene promoters were more likely to be retained following sorbitol treatment, and that these retained CTCF peaks could be halting loop extrusion leading to new loop formation.

Because we had previously observed phase separation-dependent nuclear localization and 3D clustering of YAP1 in response to sorbitol, we also mapped YAP1 occupancy using CUT&Tag. Surprisingly, we saw a global loss of YAP1 binding with 12,553 decreasing and only 21 increasing occupancy in response to sorbitol (DESeq2, padj < 0.05) (**Figure 3A**). YAP1 occupied many (68.3%) gained loop anchors even in untreated cells and was modestly retained at these sites following treatment (**Figure 3B**) but not to the extent of CTCF and RAD21. In contrast to CTCF and cohesin, YAP1 occupancy showed strong depletion both at and between loop anchors genome-wide (**Figure 3C**). Taken together, these results indicate that YAP1 is unlikely to be a critical driver of sorbitol-induced looping.

We also mapped the occupancy profiles of histone H3K27 acetylation (H3K27ac) as it is a known marker of active enhancers and promoters and has previously been correlated with loop formation in the absence of CTCF**^26^**. Indeed, H3K27ac was enriched at sorbitol-induced loop anchors both before and in response to sorbitol treatment (**Figure S9**); however, no change was observed in H3K27ac at gained or lost loop anchors, nor was there a change in occupancy at or between loop anchors (**Figure S9**). While we cannot rule out H3K27ac’s role in sorbitol-induced loop formation, these results do not provide strong evidence of it.

### Sorbitol-induced loop formation require cohesin but not CTCF

Due to their enrichment at sorbitol-induced loop anchors, we used perturbation experiments to directly test whether YAP1, RAD21, or CTCF were required for sorbitol-induced loop formation. First, we mapped 3D chromatin structure in cells expressing eGFP-YAP1Δ-TAD in which the intrinsically disordered transcription activation domain (TAD) of YAP1 was deleted. This has previously been shown to function as a dominant negative, inhibiting phase separation of exogenous YAP1 and preventing both nuclear localization and condensate formation**^20^**. APA analysis of the resulting Hi-C data (**Figure 4A**) revealed no impact on sorbitol-induced looping, demonstrating that YAP1 is not responsible for sorbitol-induced loop formation. To test the requirement of canonical loop extrusion machinery, we acutely depleted CTCF or RAD21 in HCT116 cells using an auxin-inducible degron system, treated cells with 200 mM sorbitol for 1 hour, and mapped 3D chromatin contact frequencies using Hi-C. Efficient depletion of both CTCF and RAD21 was confirmed by quantification of mean nuclear GFP intensity in the endogenously tagged HCT116 degron lines, with near-complete loss of nuclear signal observed for both proteins following auxin treatment **(Figure S10)**. CTCF depletion also had no clear impact on sorbitol-induced loop formation (**Figure 4A**), suggesting that it is not required for loop formation under hyperosmotic stress. In contrast, RAD21 depletion completely eliminated sorbitol-induced loop enrichment **(Figure 4A)**, demonstrating that cohesin-mediated extrusion is essential for sorbitol-induced loop formation.

**Fig. 4.**
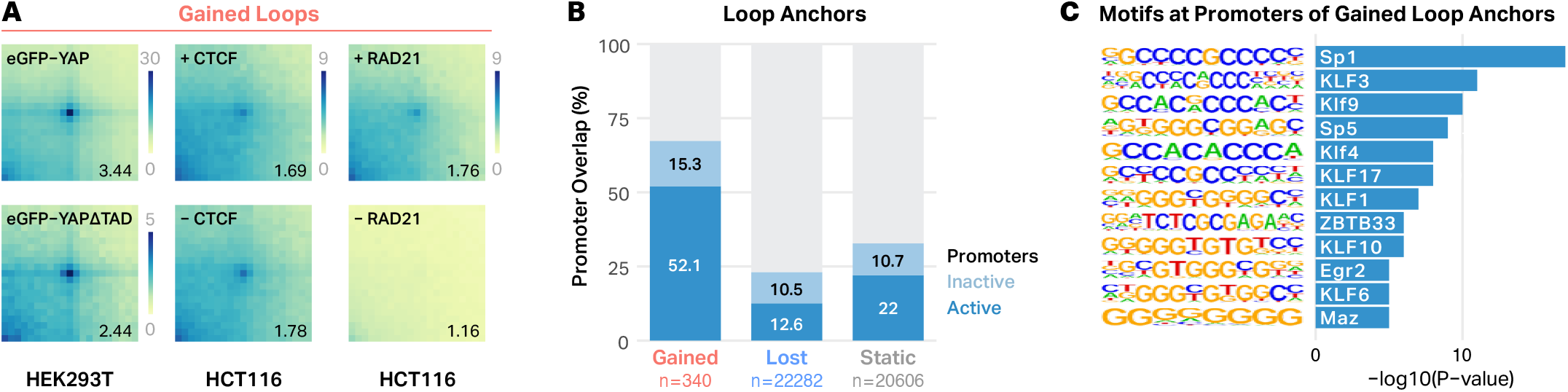
Sorbitol-induced chromatin loops require RAD21 and are enriched at promoters. **A**, Aggregate peak analysis (APA) plots showing the impact of sorbitol-induced looping after expression of a dominant negative YAP1 (eGFP-YAPdTAD) (left), degradation of CTCF (middle), or degradation of RAD21 (right). APA scores are indicated in the bottom-right corner of each matrix. Acute loss of RAD21 markedly reduces sorbitol-induced loop signal, whereas CTCF depletion and expression of the YAP1 dominant negative have no discernable impact. **B**, Proportion of gained, lost, and static loop anchors that overlap annotated promoters, stratified by active (blue) and inactive (light blue) promoter status based on RNA-seq expression at the 0h baseline (normalized count > 5). **C**, Enriched transcription factor binding motifs at promoters of gained loop anchors include SP1, KLF, and related promoter-associated transcription factors, identified by HOMER motif analysis.

### Sorbitol-induced loop anchors are enriched at gene promoters

We next asked whether alternative genomic features —particularly regulatory elements—might preferentially anchor sorbitol-induced loops. Interestingly, sorbitol-induced loop anchors displayed a strong promoter bias: 52.1% overlapped annotated promoters, compared to 12.6% of lost anchors and 22% of static anchors **(Figure 4B)**. Of these promoter-overlapping anchors, the majority corresponded to transcriptionally active promoters as defined by baseline RNA-seq expression, indicating that sorbitol-induced loops preferentially form at loci with pre-existing transcriptional activity **(Figure 4B)**. Furthermore, 53.7% of sorbitol-induced loops were classified as enhancer– promoter interactions, compared to only 16.0% of lost and 20.4% of static loops, supporting the enrichment of stress-induced loops at active regulatory interactions **(Figure S11A)**.

To test whether this regulatory-element bias reflected a broader change in cis-regulatory element (CRE) communication, we quantified interactions between putative CREs. We defined putative CREs as sites with overlapping H3K27ac and chromatin accessibility peaks. We took the 1,000 CREs with the strongest H3K27ac signal and created APA plots that aggregated contact frequency between all pairs of CREs separated by 25kb to 1Mb (n = 791 pairs; **Figure S11B)**. These plots revealed a loss of stripes, which we attribute to a global decrease in cohesin-mediated extrusion, and a slight decrease in contact frequency between putative CREs, consistent with the decrease we observe in compartmentalization (**Figure S2**).

To determine if any specific transcription factors (TFs) might explain sorbitol-induced loop formation, we performed motif enrichment analysis. Promoter regions underlying gained anchors were enriched for GC-rich promoter-associated motifs, including multiple members of the SP and KLF transcription factor families **(Figure 4C)**. These features align with prior work showing that promoter-bound transcription factors stabilize cohesin and organize long-range interactions in contexts where CTCF occupancy is reduced or insufficient for anchoring**^27–29^**. Comparative motif analysis further confirmed that SP/KLF motifs are selectively enriched at retained CTCF and RAD21 sites relative to lost sites and at sorbitol-induced loop anchors relative to lost anchors, while CTCF motif enrichment was comparable between retained and lost categories **(Figure S12A–F)**. Integration of ENCODE HEK293 ChIP-seq data showed that gained loop anchors were associated with a higher likelihood of overlap with binding sites for nearly all tested TFs **(Figure S12G–I)**. These results suggest that CTCF retention is likely associated with highly active regions rather than specifically tied to SP or KLF family TFs themselves.

Together, these findings suggest that during hyperosmotic stress, cohesin-mediated extrusion persists but is preferentially stabilized at promoter-proximal sites where CTCF is selectively retained.

### Sorbitol-induced loop formation precedes expression of anchor genes

The strong promoter bias of sorbitol-induced loop anchors **(Figure 4B)** suggested that at least some of these structural changes might participate in the transcriptional response to hyperosmotic stress. To define this response, we performed a seven-point RNA-seq time course in HEK293T cells treated with 200 mM sorbitol (0, 1, 3, 6, 9, 12, and 24 hours). Relative to untreated controls, we observed a transcriptional program in which the number of significantly differentially expressed genes (absolute log2 fold change > 2, padj < 0.05) increased from a few hundred at 1 hour to over 900 by 24 hours, with the response overwhelmingly dominated by gene activation **(Figure 5A)**. Like-lihood-ratio testing identified 6,097 genes with significant time-dependent behavior, which were clustered based on their z-scored expression profiles **(Figure 5A)**. Consistent with previous reports, we detect down-stream of gene (DoG) transcription, although the effect is modest and restricted to select loci **(Figure S13)^18,22^**. Gene Ontology (GO) enrichment analysis revealed that upregulated genes were enriched for biological processes associated with cell-cell interactions and stress signaling, including leukocyte cell-cell adhesion, regulation of inflammatory response, angiogenesis, response to mechanical stimulus, and sodium ion transport **(Figure 5B)**. Downregulated genes were not enriched for any specific biological processes.

**Fig. 5.**
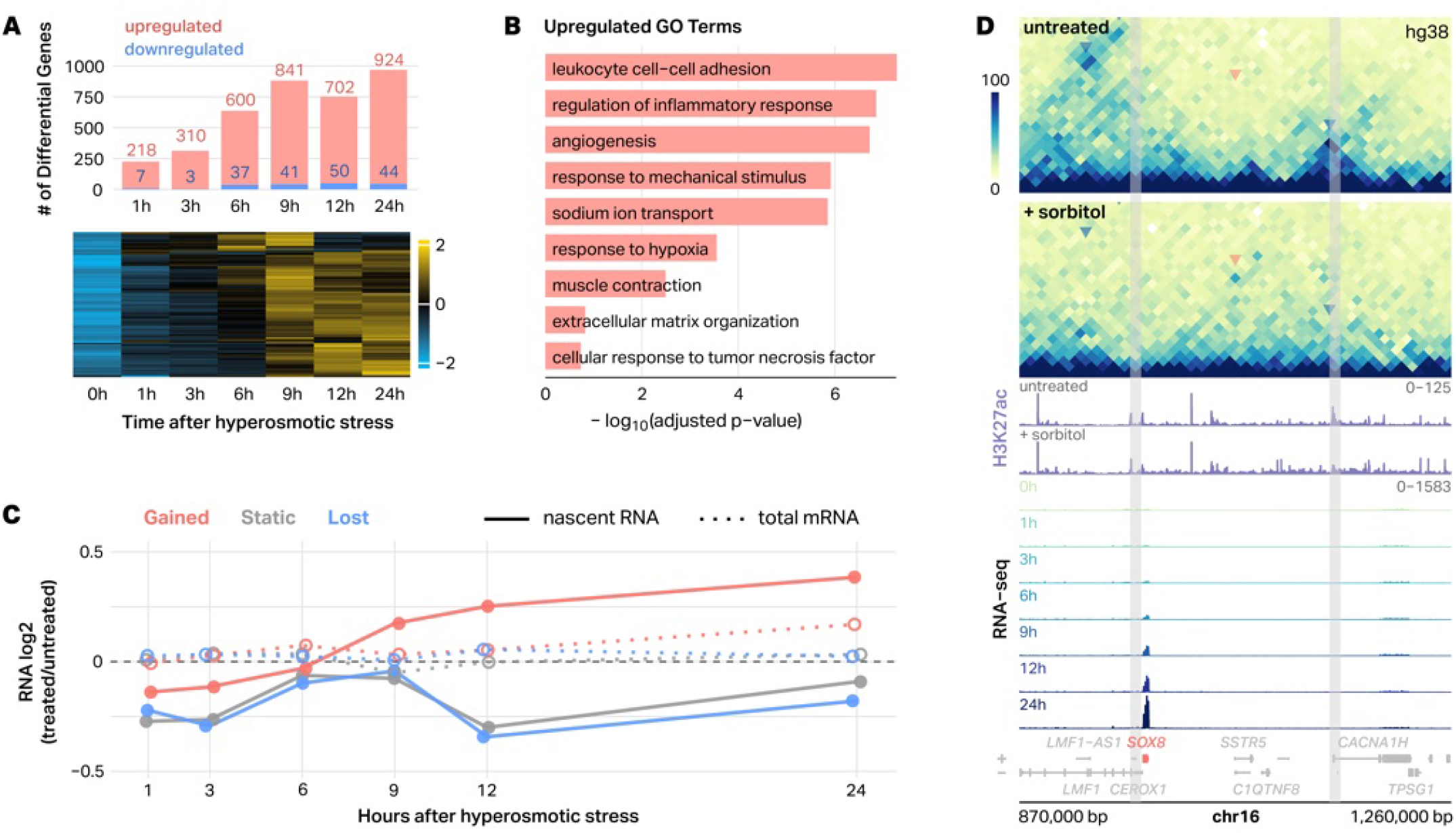
Sorbitol-induced loops are associated with transcriptional changes. **A**, Bar plot depicting differentially expressed genes (|log**2** fold-change| > 2, padj < 0.05) in HEK293T cells treated with 200 mM sorbitol with labels to indicate number of up- and downregulated genes. Heatmap shows z-scored expression of 6,097 LRT-identified genes (DESeq2, LRT, p < 0.05), revealing transient, transcriptional programs across the time course. **B**, Gene Ontology (GO) enrichment of genes significantly upregulated or downregulated at any time point. Upregulated genes are enriched for inflammatory, cell–cell adhesion, and mechanical response pathways, while downregulated genes are enriched for RNA modification and translational control processes (ranked by –log10 adjusted p-value). **C**, Median transcriptional responses of genes at gained, static, and lost loop anchors derived from Exon-Intron Split Analysis (EISA). Solid lines show nascent transcription (Δintron); dotted lines show total mRNA changes (Δexon). Promoters were assigned to loop classes based on overlap with sorbitol-induced, unchanged, or lost anchors. Lines show median RNA log**2**(sorbitol/control) across the time course. Genes at gained anchors exhibit stronger and sustained induction of nascent transcription compared with static or lost groups, with the transcriptional response emerging several hours after loop formation. Asterisks denote time points where the mean log**2**FC for gained-anchor genes differs significantly from zero (*p < 0.05, **p < 0.01, ***p < 0.001; one-sample t-test, Benjamini-Hochberg adjusted). **D**, Example chromosome 16 locus (hg38) showing a sorbitol-induced loop linked to transcriptional activation. Hi-C contact maps (10-kb) show loop gain, H3K27ac CUT&Tag tracks show cis-regulatory element activity before and after sorbitol treatment, and RNA-seq tracks (0–24 h) show progressive SOX8 activation.

These results confirm previous findings and demonstrate that hyperosmotic stress elicits a coordinated transcriptional program characterized by activation of stress-responsive and signaling-related pathways**^9,10,18^**. Moreover, only 27.2% of sorbitol-responsive genes localize to gained or lost loop anchors, whereas the remaining 72.8% fall outside these regions, indicating that loop remodeling accounts for only a subset of the transcriptional response. Genes whose promoters lie at sorbitol-induced loop anchors show increased expression in response to stress, with this upregulation emerging several hours after loop formation **(Figure 5C)**. To distinguish transcriptional from post-transcriptional contributions to this delayed response, we performed Exon-Intron Split Analysis (EISA)**^30^**, using intronic reads as a proxy for nascent transcription. Genes at gained loop anchors show a stronger and more sustained increase in nascent RNA relative to static or lost anchor genes, with the delay relative to loop formation preserved, consistent with a transcriptional rather than post-transcriptional mechanism of gene activation **(Figure 5C)**. These changes are exemplified at the SOX8 locus **(Figure 5D)**. H3K27ac CUT&Tag tracks at this locus reveal a pre-existing cis-regulatory element at the distal loop anchor, supporting the classification of this interaction as an enhancer–promoter loop **(Figure 5D)**. Hi-C maps reveal the formation of a *de novo* loop connecting the SOX8 promoter and a distal region, coinciding with a progressive increase in SOX8 expression across the 24-hour time course **(Figure 5D)**. To investigate whether cohesin-dependent sorbitol induced looping contributes to transcriptional activation at gained anchors, we examined gene expression in auxin-inducible RAD21-depleted HCT116 cells treated with sorbitol. In response to sorbitol, genes at sorbitol-induced loop anchors in cells lacking RAD21 showed a modest but significant reduction in expression relative to cells expressing RAD21 (p < 0.001, n = 405 genes), whereas genes at pre-existing loop anchors were unaffected (p = 0.423, n = 3,026 genes). These results are consistent with a role for cohesin in supporting transcriptional activation at stress-induced loop anchors, although the magnitude of the effect suggests that cohesin-dependent looping is likely only one contributor to this response. Together, these analyses demonstrate that gained loops are preferentially anchored at promoters that become transcriptionally activated during the hyperosmotic stress response, linking stress-induced 3D chromatin structure to a subset of downstream gene expression programs.

## Discussion

Here, we characterized changes to 3D chromatin structure, protein-DNA interactions, and gene expression during the response of human cells to hyperosmotic stress. Our findings both agree with and extend those of previous studies. In agreement with prior work, we see evidence for loss of CTCF-mediated looping, run-on transcription, and large-scale changes in gene expression**^18,19^**. We extend those results by providing evidence of the formation of a new class of sorbitol-induced loops. These loops exhibit slight phenotypic differences to canonical CTCF-mediated loops, which could indicate different mechanisms of formation. One intriguing feature was that sorbitol-induced loops manifested as more punctate spots in Hi-C maps. These punctate structures are reminiscent of the punctate loops observed in Drosophila, which also do not involve CTCF**^31^**. Whether or not sorbitol-induced loops form via the same mechanisms as these Drosophila loops is unclear, as is why CTCF-driven loops are less punctate. These results offer another important data point that might help resolve such mysteries. We also extended previous knowledge by providing data on the timing and persistence of sorbitol-induced loop loss and formation. The timing of lost and gained loops showed a strong anticorrelation, which suggests that both may rely on the same underlying mechanisms. Indeed, our results are consistent with the loss of CTCF binding driving both looping phenotypes; however, proving this mechanism will require further investigation. This project was largely motivated by our previous work that suggested YAP1 might mediate 3D chromatin changes in response to hyperosmotic stress; however, the results from this study only partially agree. In our previous work, we found that in response to sorbitol, YAP1 enters the nucleus and forms condensates that colocalize with hubs of open chromatin. In the current study, we did observe large-scale changes in 3D chromatin structure, including the formation of new loops whose anchors were enriched for YAP1. Surprisingly, however, the expression of a dominant negative form of YAP1, which we previously found inhibits nuclear localization and condensate formation (of both exogenous and endogenous YAP1), did not inhibit the formation of sorbitol-induced chromatin loops. This is strong evidence that YAP1 is not required for sorbitol-induced loop formation. One possible explanation for the discrepancy between our current Hi-C-based results and our previous 3D ATAC-PALM results is that 3D ATAC-PALM and Hi-C probe chromatin organization on fundamentally different spatial scales. Hi-C requires that two regions be within a crosslinking radius, which some estimate to be as small as tens of nanometers**^32–35^**. In contrast, even with very high resolution analysis, 3D ATAC-PALM can only resolve colocalization events down to a radius of 100 to 400nm**^20,36^**. Therefore, the type of clustering observed by 3D ATAC-PALM may not be sufficient for detection by Hi-C. Further work will be required to disentangle the precise role of YAP1 in sorbitol-induced 3D chromatin changes, but the existing data are consistent with YAP1 playing a role in large-scale changes to nuclear localization but no role in changes to chromatin looping.

The enrichment of active gene promoters, enhancers, and GC-rich transcription factor motifs at sorbitol-induced loop anchors is consistent with a number of studies showing that active regulatory loci can stabilize 3D chromatin contacts, sometimes independently of CTCF. Cohesin degradation has been shown to induce interactions between super enhancers. Multiple studies have implicated SP and KLF family TFs, whose motifs were enriched at our sorbitol-induced loop anchors, in chromatin looping. However, other data argues against a simple model in which SP/KLF factors alone facilitate the formation of these loops. In particular, retained CTCF and RAD21 sites, as well as sorbitol-induced loop anchors, show broad enrichment for binding sites of most transcription factors, suggesting that SP/KLF motifs may simply mark highly accessible, transcriptionally active promoters rather than serve as the primary drivers of sorbitol-induced loop formation. Thus, we favor a model in which active promoter machinery— potentially including SP/KLF-family factors, other promoter-associated transcription factors, RNA polymerase II, chromatin regulators, or combinations of these features—creates a chromatin environment that preferentially stabilizes cohesin during hyperosmotic stress. Direct perturbation of individual SP/KLF factors or their binding motifs will be needed to test whether they contribute causally to loop formation, although such experiments may be complicated by the extensive redundancy among SP and KLF family members and by the broader role of GC-rich promoter elements in maintaining promoter accessibility and activity.

Since sorbitol-induced loops preferentially form at transcriptionally active promoters rather than merely annotated promoters, it is possible that promoter activity itself may contribute to loop formation. Promoter-associated factors, including RNA Pol II, may help stabilize CTCF and cohesin at these sites during stress; however, because loop formation precedes the major transcriptional response, our data favor a model in which promoter-associated factors facilitate cohes-in-dependent loop formation rather than transcription itself generating these interactions. Future studies directly perturbing RNA Pol II activity during hyperosmotic stress will be needed to test this possibility.

Genes at the anchors of sorbitol-induced loops exhibit a modest but significant increase in expression in response to sorbitol. Importantly, both total and nascent transcription do not significantly increase until several hours after loop formation. In fact, gene expression peaks at 24 hours, at which time the sorbitol-induced loops have largely disappeared. This timing argues against a model in which loop formation is merely a passive consequence of transcriptional activation. At the same time, our data do not allow us to conclude that the loops directly cause the subsequent increase in gene expression. Rather, loop formation and transcriptional activation may represent parallel consequences of the hyperosmotic stress response or reflect a more complex regulatory relationship. It is important to note that the changes in gene expression at sorbitol-induced loop anchors are modest and variable. Moreover, the majority of sorbitol-responsive genes are not found at gained or lost loop anchors. Taken together, these findings suggest that chromatin loops are just one of many regulatory mechanisms that contribute to the transcriptional response to hyperosmotic stress.

There are several limitations to this study. First, our conclusions are derived from bulk Hi-C, CUT&Tag, and RNA-seq data, which average over heterogeneous cell populations. Individual cells likely exhibit varied degrees and trajectories of chromatin collapse, loop gain, and transcriptional response that are not resolved in our current datasets. Second, crosslinking-based assays provide static snapshots of nuclear organization, limiting our ability to infer causality or fine-grained temporal order between specific looping events and changes in transcription beyond the coarse time resolution of our time courses. Third, our mechanistic perturbations rely on engineered cell lines (eGFP-YAP1 and eGFP-YAP1ΔTAD HEK293T cells; auxin-inducible CTCF and RAD21 degrons in HCT116 cells), which may differ in chromatin context and osmoadaptive pathways from primary renal, immune, or other physiologically relevant cell types. Finally, we focused on a subset of architectural and regulatory factors; additional contributors—including other cohesin regulators, chromatin remodelers, or nuclear envelope components—may participate in shaping the stress-induced 3D genome but were not directly examined here.

Together, our results reveal that hyperosmotic stress triggers a rapid, global, and reversible rewiring of 3D chromatin structure characterized by the collapse of canonical CTCF-mediated loops and the emergence of a distinct class of CTCF-independent, cohesin-dependent sorbitol-induced loops. These findings demonstrate how physical genome organization adapts under acute environmental stress and support accessible promoters as alternative architectural scaffolds during periods of widespread chromatin destabilization. More broadly, this study highlights the remarkable plasticity of the 3D genome and suggests that environmental stress can transiently reshape nuclear architecture through mechanisms that are fundamentally distinct from those operating during homeostasis.

**Acknowledgements**

We thank Erika Deoudes for data visualization, illustration, proofreading, and typesetting. We thank Samantha Pattenden for use of the Covaris LE220 instrument, which was provided by the North Carolina Biotechnology Center Institutional Development Program grant 2017-IDG-1005. We also thank Brian Golitz and the UNC CRISPR Core for technical assistance. We thank Wendy Salmon and the UNC Hooker Imaging Core for assistance with fluorescence microscopy, including use of the GE IN Cell Analyzer 2200 and the Leica Stellaris 8 FALCON STED confocal microscope (NIH grant 1S10OD030300).

## Data Availability

All raw and processed sequencing data generated in this study have been submitted to the NCBI Gene Expression Omnibus (GEO; https://www.ncbi.nlm.nih.gov/geo/). The Hi-C data are available under accession number GSE310051. The Hi-C data for HCT116-RAD21-mAID2 and HCT116-CTCF-mAID2 cells are available under accession number GSE312288. The RNA-seq data are available under accession number GSE310049. The RNA-seq data for HCT116-RAD21-mAID2 cells are available under accession number GSE335237. The CUT&Tag data are available under accession number GSE310047. The ATAC-seq data available under accession number GSE329313. The code to process and analyze these data is available on GitHub (https://github.com/jpflores-13/STRS).

## Author Contributions

J.P.F. conceived the study, designed experiments, performed analyses, curated data, and wrote the original draft. A.A.P., Z.D., C.X., D.C.A., G.P., Y.W., I.Y.Q.-B., H.K. T.R., J.D.B., G.C.F., Y.J.L., E.B., Y.L., and I.S. contributed to experiments, data generation, and validation. J.D., D.C., B.D.S., J.M.D., and G.W. provided methodological expertise, resources, and supervision. D.H.P. provided conceptual guidance, supervision, funding acquisition, and project administration, and served as senior author. All authors reviewed and edited the manuscript.

## Experimental model and subject details

### Cell Lines

HEK293T eGFP-YAP1, HEK293T eGFP-YAP1dTAD (gifts from the D. Cai Lab) and HEK293T (ATCC CRL-3216) cells were cultured at 37 °C and 5% CO2 in DMEM supplemented with 10% fetal bovine serum (FBS; Gibco) and 100 U ml-1 penicillin/streptomycin (Fisher Scientific). HEK293T and its derivatives are of female origin (RRID:CVCL_0063). HCT116 (ATCC CCL-247), HCT116-RAD21-mAID2, and HCT116-CTCF-mAID2 (obtained from M. Kanemaki Lab) cells were cultured at 37 °C and 5% CO2 in McCoy’s 5A medium supplemented with 10% FBS (Gibco) and 100 U ml-1 penicillin/ streptomycin (Fisher Scientific). HCT116 and its derivatives are of male origin (RRID:CVCL_0291). Unless noted, cells were grown on 10-cm Corning plates. Cells were checked for mycoplasma using Genetica Cell Line Testing (LabCorp) and confirmed negative. Parental lines were obtained directly from ATCC (authenticated at deposit), and engineered derivatives were used as received from the originating labs.

## METHODS

### Sorbitol Treatments

For high-depth Hi-C experiments, wild-type HEK293T, eGFP-YAP1 HEK293T, and wild-type HCT116 cells were serum-starved for 1 h and treated with control serum-free media or 0.2 M sorbitol (MP Biomedicals, cat #194742) in serum-free media for 1 h, as described by Hong et al**^37^**. For Hi-C time course experiments, wild-type HEK293T cells were serum-starved for 1 h and treated with control serum-free media for 1 h or 0.2 M sorbitol for 10 m, 30 m, 1 h, 3 h, 6 h, 12 h, and 24 h. For low-depth Hi-C experiments, HEK293T eGFP-YAP1dTAD cells were serum-starved for 1 h and treated with 0.2 M sorbitol for 1 h. For RNA-seq time course experiments, wild-type HEK293T cells were serum-starved for 1 h and treated with control serum-free media for 1 h or 0.2 M sorbitol for 1 h, 3 h, 6 h, 9 h, 12 h, and 24 h.

### Auxin Treatments

HCT116-RAD21-mAID2 and HCT116-CTCF-mAID2 cells were serum-starved and treated with 1 µM 5-phenyl-indole-3-acetic acid (5-Ph-IAA) (BioAcademia, Japan, #30-003) for 6 h, adapted from Yesbolatova et al**^38^**. Cells were then treated with 0.2 M sorbitol in serum-free media with or without 1 µM auxin (5-Ph-IAA) for 1 h. For RNA-seq after auxin-mediated cohesin degradation, HCT116 mAID-RAD21 cells were serum-starved for 1 h and treated with control serum-free media for 6 h, 0.2 M sorbitol for 6 h, or 0.2 M sorbitol and 1 µM 5-Ph-IAA for 6 h.

### Fluorescence Microscopy

For immunofluorescence imaging of CTCF and RAD21 protein depletion, HCT116-CTCF-mAID2 and HCT116-RAD21-mAID2 cells were seeded in 6-well plates and treated with 1 µM 5-phenylindole-3-acetic acid (5-Ph-IAA) for 6 hours to induce auxin-dependent protein degradation, with or without subsequent 200 mM sorbitol treatment for 1 hour as indicated. Following treatment, media was removed and cells were washed gently once with PBS. Cells were fixed in 4% paraformaldehyde (Fisher Scientific, Cat# 28909) for 10–15 minutes at room temperature, then washed 2–3 times with PBS. Nuclei were stained with Hoechst 33342 (1:10,000 dilution in PBS) for 5–10 minutes at room temperature protected from light, followed by a final wash with PBS. Cells were imaged in PBS. High-content widefield fluorescence imaging was performed on a GE IN Cell Analyzer 2200 at the UNC Hooker Imaging Core. Confocal imaging was performed on a Leica Stellaris 8 FALCON STED laser scanning confocal microscope at the UNC Hooker Imaging Core (supported by NIH grant 1S10OD030300). Imaging assistance was provided by Wendy Salmon. Image analysis was performed using CellProfiler (v4.2.8).

### Live-cell imaging of CTCF and RAD21 redistribution

HEK293T cells were seeded in 8-well chambered coverglass plates designed for high-resolution live-cell imaging (Cellvis, C8-1.5P) and transfected with either mEmerald-RAD21-N-18 (Addgene plasmid #54248; RRID:Addgene_54248) or pKS070-pCAGGS-3XFLAG-(human)CTCF-eGFP (Addgene plasmid #156448; RRID:Addgene_156448) using Lipofectamine 3000 (Invitrogen, L3000015). Cells were incubated for 24 h at 37 °C in DMEM before the culture medium was replaced with FluoroBrite DMEM (Gibco, A18967-010) to reduce background fluorescence during live-cell imaging. Nuclei were stained with 4 μM Hoechst 33342 (Thermo Fisher Scientific, 62249) in PBS for 10 min. Cells were then treated with either FluoroBrite DMEM alone (control) or 0.2 M sorbitol (Sigma, 102111476) prepared in FluoroBrite DMEM for 1 h. Live-cell fluorescence imaging was performed using a Zeiss LSM 900 confocal microscope equipped with Airyscan 2 to visualize the nuclear distribution of CTCF-EGFP and RAD21-mEmerald.

### Nuclear volume quantification

HEK293T cells were seeded onto 24-well glass-bottom plates (Cellvis, P24-1.5H-N) and grown to approximately 70% confluency. Prior to imaging, cells were stained with 0.2 μM Hoechst 33342 (Thermo Fisher Scientific, 62249) for 3 min and washed once with imaging medium consisting of FluoroBrite DMEM (Gibco, A1896702) supplemented with 5% FBS. Sorbitol (Sigma-Aldrich, S1876) was dissolved in imaging medium and added to a final concentration of 0.2 M. Live-cell imaging was performed using a Zeiss LSM 900 confocal microscope equipped with a 20× objective. Ten fields of view were selected and imaged immediately before treatment and again after 1 h of sorbitol exposure. Images were acquired using a 647 × 647 pixel field of view, 4× zoom, scan speed 8, averaging 2, laser power 0.5%, and detector gain 600. Three-dimensional image stacks were collected at 0.3 μm intervals across 50 optical sections centered on the nucleus. Nuclear segmentation and volume measurements were performed using CellProfiler (v4) with a custom analysis pipeline. A total of 416 control nuclei and 436 sorbitol-treated nuclei were analyzed. Statistical significance was assessed using a two-tailed Welch’s t-test.

### Hi-C Library Generation

Cells were treated in serum-free, sorbitol-containing DMEM as described above, then crosslinked in 1% formaldehyde (Fisher Scientific, Cat# 28908) for 10 min at room temperature and quenched with 0.2 M glycine (Fisher Scientific, Cat# 07-678-003) for 5 min. Cells were washed twice with ice-cold PBS, pelleted (∼5 × 106 cells/ pellet), snap-frozen in liquid nitrogen, and stored at -80 °C. In situ Hi-C libraries were prepared following the protocol of Rao et al. 2014**^35^**, including nuclei isolation, MboI digestion, biotin-dATP fill-in, proximity ligation, crosslink reversal, and DNA purification**^35^**. Genomic DNA was sheared to 300–500 bp using a Covaris LE220 (DF 20, PIP 100, 200 cycles/burst, 80 s), followed by AMPure XP size selection. Biotinylated fragments were captured with streptavidin beads, and all subsequent steps (end-repair, A-tailing, and adaptor ligation) were performed on-bead using TruSeq Nano adapters. Libraries were amplified by PCR (7–10 cycles), quality-checked (Qubit dsDNA HS and Agilent TapeStation D1000), pooled to a final concentration of 10 nM, and sequenced in paired-end 150 bp mode on an Illumina NovaSeq X Plus using an S4 flow cell (2 × 150 bp).

### RNA-seq Library Generation

Total RNA was extracted using the QIAGEN RNeasy Mini Kit (Cat. #74104/74106) with on-column RNase-free DNase I (Cat. #79254) per manufacturer’s protocol. RNA integrity was assessed by Qubit and Agilent TapeStation. Extracted RNA was shipped on dry ice to Azenta Life Sciences, where total RNA rRNA depletion and stranded RNA-seq library preparation (KAPA RNA HyperPrep with RiboErase HMR) were performed. Libraries were sequenced paired-end 150 bp on an Illumina NovaSeq X-series instrument, targeting ∼30 million read pairs per sample.

### CUT&Tag Library Generation

CUT&Tag was performed as described by Kaya-Okur et al. with minor modifications**^39^**. Briefly, ∼1 × 105 cells per sample were harvested and supplemented with 5% GFP-H2B-expressing NIH3T3 cells for spike-in normalization. Cells were washed in Wash Buffer (HEPES, NaCl, spermidine, protease inhibitors) and bound to concanavalin-A-coated magnetic beads (10 µL/sample; activated in Bead Activation Buffer) for 10 min at room temperature (RT). Bead-bound cells were resuspended in Digitonin 150 Buffer (±2 mM EDTA) and incubated with primary antibodies overnight at 4 °C. Primary antibodies were used at 1:50 and included anti-GFP to target eGFP-YAP1 (Abcam, #ab290), anti-CTCF (Cell Signaling Technology, #3418S), anti-RAD21 (Abcam, #ab217678), and anti-H3K27ac (Abcam, #ab4729). After washing, cells were incubated with secondary antibody (1:50, 30 min, RT), followed by the pre-loaded pA-Tn5 adapter complex (1:200 in Digitonin 300 Buffer) for 1 h at RT. Unbound enzyme was removed and tagmentation was initiated in Digitonin 300 Buffer supplemented with 10 mM MgCl2 (37 °C, 1 h). DNA was released with SDS Release Buffer (58 °C, 1 h), quenched with SDS Quench Buffer, and purified using AMPure XP beads. Indexed libraries were generated using universal i5 and barcoded i7 primers, PCR-amplified with 2× master mix, cleaned with 0.9× AMPure XP, eluted in 10 mM Tris-HCl (pH 8.0), and sequenced (PE75) on an Illumina NextSeq 500 or equivalent platform.

### ATAC-seq Library Generation

ATAC-seq libraries were prepared following the Omni-ATAC protocol as described in Corces et al.**^40^**. Briefly, eGFP-YAP1 HEK293T cells were serum-starved for 1 hour and treated with vehicle (serum-free media) or 200 mM sorbitol for 1 hour prior to harvest. 50,000 cells per sample were pelleted at 500 × g for 5 minutes at 4 °C. Cell pellets were resuspended in cold ATAC lysis buffer (10 mM Tris-HCl pH 7.4, 10 mM NaCl, 3 mM MgCl2, 0.1% NP-40, 0.1% Tween-20, 0.01% digitonin) and incubated on ice for 3 minutes. Nuclei were pelleted by centrifugation at 500 × g for 10 minutes at 4 °C, washed once with wash buffer (10 mM Tris-HCl pH 7.4, 10 mM NaCl, 3 mM MgCl2, 0.1% Tween-20), and resuspended in 50 µL of transposition mix containing Nextera XT transposase (Illumina). Tagmentation was carried out at 37 °C for 30 minutes with gentle shaking. Transposed

DNA was purified using the Zymo DNA Clean and Concentrator kit. Libraries were amplified using NEBNext High-Fidelity 2× PCR Master Mix (NEB, cat# M0541L) for 5 initial cycles, followed by a quantitative PCR side reaction on 5% of the amplified material to determine the number of additional cycles required (typically 4–7 additional cycles). Libraries were cleaned up using a two-sided AMPure XP bead selection (0.5× then 1.3×), quantified by Qubit (dsDNA HS Assay) and the KAPA Library Quantification kit, and assessed for quality by Agilent TapeStation (D1000). Libraries were pooled at equimolar concentrations and sequenced in paired-end 75 bp mode on an Illumina NextSeq 500.

## Quantification and statistical analysis

### Hi-C Processing and Loop Detection

Hi-C data were processed using a custom pipeline called *dietJuicer* (Snakemake workflow; https://github.com/PhanstielLab/dietJuicer), derived from Juicer**^41^**. Reads were aligned to the hg38 genome using bwa mem (v0.7.17) and filtered to valid pairs (MAPQ ≥ 30)**^42^**. In situ Hi-C libraries were generated using MboI restriction digestion**^35^**. Hi-C contact matrices were generated at 5-kb and 10-kb resolutions and balanced using SCALE normalization for visualization and APA plots. Chromatin loops were identified using SIP (v1.6.1) (https://github.com/PouletAxel/SIP) with parameters `-g 1 -t 2000 -fdr 0.05`(using `-isDroso false`), and loop anchors were lifted to a 10-kb grid and merged across replicates using DBSCAN clustering (ε = 20 kb; Manhattan distance)**^31,43,44^**. For differential loop analysis, raw observed loop pixel counts were extracted at 10-kb resolution directly from per-sample `.hic` files using strawr (v0.0.9) as implemented in mariner, ensuring no matrix normalization was applied (i.e., `norm = NONE, matrix = observed`)**^45^**. SCALE-normalized matrices were used exclusively for visualization and aggregate peak analysis.

### Chromatin Compartment and Domain Analysis

Chromatin A/B compartments were called from ICE-balanced Hi-C contact matrices at 10-kb resolution using the leading eigenvector (PC1) of the intrachromosomal correlation matrix (‘cooltools eigscis’**^46^**). PC1 sign was oriented per chromosome using a GC-content phasing track such that positive values correspond to the A compartment and negative values to the B compartment. Compartment strength was quantified from saddle plots (‘cooltools saddle‘**^46^**, 50 bins) generated from the control-sample eigenvector and condition-matched expected contact frequency, and summarized per replicate as (AA + BB) / (AB + BA) using the three corner bins of the saddle matrix. Chromatin domain boundaries were identified using a diamond insulation score with a 250-kb sliding window (‘cooltools insulation’**^46^**) on the same balanced matrices; mean insulation profiles were computed across called boundaries (+/-500 kb) to compare boundary strength between conditions and genotypes.

### Differential Loop Analysis

Loop counts were modeled in DESeq2 with the design `∼ Replicate + Treatment`**^47^**. Because hyperosmotic stress induces global changes in chromatin interaction frequencies, we expected widespread differences in total loop counts between conditions. To avoid normalization procedures that assume global count stability and could therefore mask true genome-wide remodeling, size factors were fixed to 1 (`sizeFactors(dds) <-1`), ensuring loops were compared on an unscaled basis. Wald tests were used for contrasts, and log2 fold-changes were shrunk using apeglm**^48^**. Unless stated otherwise, gained loops were defined as padj < 0.1 with log2FC > 0, and lost loops as padj < 0.1 with log2FC < 0. For QC and visualization, variance-stabilized counts were converted to Z-scores and clustered by k-means.

### Aggregate Hi-C analysis

Aggregate loop and domain analyses were performed using mariner (v1.5.0) and strawr (v0.0.9). KR-or VC_ SQRT-normalized Hi-C contact matrices were extracted at 10-kb resolution. Loops with inter-chromosomal anchors or insufficient signal (row/column sums < 1) were excluded. Loop-centric aggregates were generated by centering windows on loop pixels with a ±0.5-loop-width buffer. For domain-centric aggregates, windows were scaled to loop span and subsequently resized to a 100 × 100 matrix using bilinear interpolation. Each window was normalized by total contact count prior to averaging across loops.

Loop strength was quantified as enrichment of the central pixel (or central 3×3 window) relative to a local background, defined either as the median of flanking ring bins or corner regions (for APA-style summaries). One-dimensional enrichment profiles were derived by extracting diagonal traces and normalizing to the corresponding background estimate.

For comparisons to size- and contact-matched null loop sets, we used the matchRanges function from the nulllranges package (v1.14.0) to select control loops matched on log(loop size) and log(aggregated contact frequency) using stratified sampling without replacement.

### Hi-C time course analysis

We analyzed chromatin loop dynamics in wild-type HEK293T cells (0 h, 10 min, 30 min, 1 h, 3 h, 6 h, 12 h, 24 h; control and sorbitol). Differentially gained and lost loops (DESeq2) were used as input, and subsampled Hi-C maps were processed in R with mariner (v1.5.0) and visualized with plotgardener (v1.5.0)**^45,49^**.

Aggregate peak analysis (APA) was performed at 10-kb resolution using ±10-bin windows, with observed counts extracted by `pullHicMatrices` and normalized by loop number. Loop enrichment was calculated with mariner’s `calcLoopEnrichment`, which defines the foreground as the loop center and background as the top-left and bottom-right corners. Enrichment was computed as median(foreground + 1)/median(background + 1), controlling for distance-dependent decay.

### Matched chromatin loop sets

To compare gained loops against size/contact-matched controls, we used the matchRanges function from the nullranges package (v1.14.0) (stratified, no replacement) with covariates log(loop size) and log(aggregated contact frequency). Aggregated contact frequency combined sorbitol and control counts per loop; zero values were treated as missing prior to log transformation.

#### CRE-CRE contact frequency analysis

To assess whether contact frequency among putative cis-regulatory elements (CREs) changed globally upon sorbitol treatment, CREs were defined as H3K27ac CUT&Tag peaks (from differential H3K27ac DESeq2 results) that overlapped an ATAC-seq peak. Peaks with a mean normalized count (baseMean) below 50 were excluded, and the 1,000 highest-confidence CREs by baseMean were retained. All intra-chromosomal pair-wise combinations of these CREs separated by 25 kb–1 Mb were enumerated (n = 791 pairs) and snapped to a 10-kb bin grid using mariner (v1.5.0). For each pair, a ±10-bin (100-kb) window was extracted from per-condition (untreated and sorbitol-treated) .hic files using pullHicMatrices (norm = NONE, matrix = observed), and matrices were aggregated by summing across all pairs and dividing by the number of pairs to generate the aggregate peak analysis (APA) plots shown in Figure S11B. APA scores were calculated as the ratio of the median value of the center pixel to the median value of 5×5-pixel corner regions (background), with a pseudocount of 1 added to both foreground and background prior to taking the ratio.

#### Motif Enrichment Analysis

Transcription factor motif enrichment was performed using HOMER**^50^** (v5.1) findMotifsGenome.pl against the hg38 genome with repeat masking (-mask) and -size given. For gained loop anchor promoters, the focus set comprised promoter windows overlapping gained loop anchors, and an explicit, expression-matched background set (promoters not overlapping gained anchors, selected via matchRanges/nullranges) was supplied with -bg rather than using HOMER’s default randomly generated genomic background. Analogous focus/background comparisons: retained versus lost CTCF peaks, retained versus lost RAD21 peaks, and gained versus lost ATAC-seq loop-anchor peaks were run identically.

#### CTCF Motif Position Weight Matrix (PWM) Scoring

CTCF peaks called by DESeq2 were classified as retained (padj ≥ 0.1) or lost (padj < 0.1, log2FC < 0) following sorbitol treatment. For each peak, a 50-bp window centered on the MACS2-called summit was extracted from the hg38 genome sequence (‘BSgenome.Hsapiens.UCSC.hg38’) and scored against the JASPAR CORE CTCF position frequency matrix (JASPAR2020, matrix MA0139.1), converted to a log2-probability-ratio PWM (pseudocount = 0.8) using TFBSTools51. Motif matches were identified with searchSeq() (unstranded, minimum relative score 60%; windows with no match at this threshold were rescored without a minimum-score cutoff), and the maximum PWM score per window was retained. PWM score distributions were compared between retained and lost peaks using a two-sided Wilcoxon rank-sum test, with effect size reported as Cohen’s d.

#### Gene Ontology (GO) Enrichment Analysis

GO enrichment (Biological Process) was performed using ‘clusterProfiler::enrichGO’ (‘org.Hs.eg.db’) separately on genes significantly up-or down-regulated (padj < 0.05, |log2FC| > 2) at any timepoint of the LRT timecourse model, using all expressed genes in the dataset as the background gene universe. Terms were retained at p < 0.05 (Benjamini-Hochberg adjusted, q < 0.2) with gene set sizes between 10 and 500 genes; the top five terms per direction, ranked by adjusted p-value, are shown.

#### Promoter and enhancer annotation

Gene promoters were defined using ‘TxDb.Hsapiens. UCSC.hg38.knownGene’, taking the default 2,200-bp window flanking each annotated TSS (2,000 bp upstream, 200 bp downstream; GenomicFeatures::promoters()). Loop anchors, CTCF/RAD21 peaks, and ATAC peaks were classified as promoter-overlapping if they intersected this window. Putative enhancers were defined as consensus H3K27ac CUT&Tag peaks that did not overlap an annotated promoter (‘GenomicRanges::setdiff’). Loops were classified as enhancer–promoter (E–P) interactions if one anchor overlapped a promoter and the other overlapped a non-promoter H3K27ac (enhancer) peak.

### RNA-seq processing and quantification

RNA-seq reads were processed using the bagPipes Snakemake workflow (https://github.com/PhanstielLab/bagPipes/tree/v2.0.0). Adapter trimming was performed with Trim Galore, and read quality was assessed using FastQC and MultiQC**^52–54^**. Transcript quantification was carried out using Salmon (v1.10.0; `--validateMappings --seqBias –gcBias --posBias`) against GENCODE hg38 reference indices**^55^**. Quantification files were imported into R using tximeta, and counts were summarized to the gene level with tximport**^56,57^**. Genes with <10 counts in fewer than 4 samples were excluded. All samples showed RIN > 9.8 and passed quality control.

### Differential gene analysis

Transcript-level Salmon**^55^** quantifications were imported with tximeta**^57^** and summarized to gene-level counts. Genes with low expression were excluded by requiring at least 50 raw counts in ≥ 4 samples prior to modeling. Gene-level counts were analyzed in DESeq2**^47^** with the design `∼ Bio_Rep + Time`, treating biological replicate and timepoint as fixed effects. To identify genes whose expression changed over the time course, we fit a likelihood ratio test (LRT) model comparing the full design (`∼ Bio_Rep + Time`) to a reduced model lacking the time term (`∼ Bio_Rep`), and defined genes with FDR-adjusted p < 0.05 as exhibiting significant time-dependent variation. Timepoint-specific responses were then quantified by extracting the coefficients for each Time_Xh_vs_0h comparison from the same model; unless otherwise stated, genes were classified as differentially expressed at a given timepoint if padj < 0.05 and |log2FC| > 2 relative to 0h. For downstream visualization, variance-stabilized counts were averaged across biological replicates, Z-scored per gene, and subjected to k-means clustering (k = 3) to distinguish early/transient from late/sustained transcriptional programs. The stacked barplot in Figure 5 reports the number of upregulated (log2FC > 2) and downregulated (log2FC < -2) genes at each timepoint, and the accompanying heatmap displays the clustered, Z-scored temporal expression profiles of these genes over the 0–24 h sorbitol time course.

### Exon-Intron Split Analysis (EISA)

To distinguish transcriptional from post-transcriptional contributions to gene expression changes, we performed Exon-Intron Split Analysis (EISA)**^30^**. RNA-seq reads were aligned to hg38 with HISAT2 and quantified with featureCounts separately for exonic and intronic regions, restricted to genes with non-overlapping annotation (GENCODE v49) to avoid ambiguous read assignment; exons were extended by 10 bp on each side before intronic regions were defined as the remaining gene body (gene body minus extended exons), following Gaidatzis et al.**^30^**. Exonic and intronic counts were modeled independently in DESeq2 with the design ∼ Bio_Rep + Time, yielding Δexon (total mRNA) and Δintron (nascent transcription) log2 fold-changes per timepoint relative to 0 h; the post-transcriptional component was calculated as Δexon – Δintron. Genes were assigned to gained, lost, or static loop classes by promoter overlap with the corresponding differential loop anchors.

### CUT&Tag processing and peak calling

CUT&Tag sequencing reads were processed using a Snakemake-based ChIP/CUT&Tag workflow derived from bagPipes (https://github.com/PhanstielLab/bag-Pipes/tree/v2.0.0). Adapter trimming and initial quality assessment were performed using Trim Galore and FastQC**^52,53^**. Reads were aligned to the hg38 reference genome using BWA-MEM (v0.7.17), and resulting alignments were sorted and indexed using samtools**^42,58^**. PCR duplicates were removed using Picard *MarkDuplicates* (`REMOVE_DUPLICATES=true`), and reads overlapping ENCODE hg38 blacklist regions were removed using bedtools (v2.31.1)**^59,60^**.

For peak calling, replicate BAM files were merged by condition and peaks were identified using MACS2 (v2.2.6; `callpeak -f BAMPE -g hs --keep-dup 1 -q 0.01`)**^61^**. Peaks were annotated using HOMER**^50^**. A unified peak set was generated by merging peak coordinates across samples using bedtools merge, and fragment counts per sample were quantified using bedtools multicov to generate a peak-by-sample count matrix for differential analysis.

Spike-in normalization was evaluated at the analysis stage. In experiments including NIH3T3 spike-in nuclei, spike-in-derived fragments were used to verify normalization stability across samples; however, no spike-in scaling factor was applied directly to read counts.

### Differential CUT&Tag Analysis

Differential peak analysis was performed separately for CTCF, RAD21, H3K27ac, and YAP1. CUT&Tag data-sets using DESeq2 with the design `∼ Replicate + Treatment`. Because hyperosmotic stress alters global chromatin occupancy, we expected widespread differences in total peak counts between conditions; to avoid normalization procedures that assume global count stability and could obscure true genome-wide changes, size factors were fixed to 1 (`sizeFactors(dds) <-1`), and peaks were compared on an unscaled basis. Peaks with low counts were removed prior to modeling by requiring a mean normalized count ≥ 15 across all samples. Wald tests were used to assess treatment effects, and log2 fold-changes were shrunk using apeglm. For CTCF and YAP1, increased peaks were defined as padj < 0.05 with log2FC > 1 and decreased peaks as padj < 0.05 with log2FC < -1. For RAD21, a slightly relaxed significance threshold (padj < 0.1) was used to account for lower signal-to-noise ratios observed in this dataset. For QC and visualization, variance-stabilizing or rlog-transformed counts were used for principal component analysis and sample-sample correlation heatmaps, and MA plots were inspected to confirm model behavior.

### ATAC-seq Processing and Peak Calling

ATAC-seq reads were processed using the bagPipes Snakemake workflow (ATACpipe.snakefile; https://github.com/PhanstielLab/bagPipes). Adapter trimming and initial quality assessment were performed using Trim Galore (v0.6.10) and FastQC (v0.12.1), with run-level summaries generated by MultiQC (v1.28). Trimmed reads were aligned to the hg38 reference genome using BWA-MEM (v0.7.17) and resulting alignments were sorted using samtools (v1.22). PCR duplicates were removed using Picard MarkDuplicates (REMOVE_DUPLICATES=true), and mitochondrial reads were excluded using samtools idxstats and view. Per-replicate peaks were called from deduplicated, filtered BAM files using MACS2 (v2.2.9.1; callpeak -f BAM -q 0.01 -g hs --nomodel --shift 100 --extsize 200 --keep-dup all -B --SPMR). Peaks from all replicates were merged across conditions using bedtools merge (v2.31.1) to generate a unified peak set. Fragment counts per peak per replicate were quantified using bedtools multicov. Per-sample signal tracks were generated using deepTools bamCoverage (v3.5.4).

### Differential ATAC-seq Analysis

Differential chromatin accessibility analysis was performed using DESeq2 (v1.44.0) on the merged peak-by-sample count matrix. The model design was specified as ∼ Replicate + Treatment to account for replicate effects while testing for treatment-dependent accessibility changes. Because hyperosmotic stress is expected to induce global changes in chromatin accessibility, size factors were fixed to 1 (sizeFactors(dds) <-1) to avoid normalization procedures that assume global count stability and could mask true genome-wide remodeling. Peaks with low counts were removed prior to modeling by requiring a mean normalized count ≥ 10 across all samples. Wald tests were used to assess treatment effects, and log2 fold-changes were shrunk using apeglm. Gained (increased) accessibility peaks were defined as padj < 0.05 with log2FC > 1 and lost (decreased) accessibility peaks as padj < 0.05 with log2FC < -1.

### Re-analysis of Publicly Available Data

Public Hi-C data from Amat et al. 2019 were downloaded from NCBI Gene Expression Omnibus (GEO) under accession number GSE111904 and processed through the same dietJuicer in-house Snakemake workflow against hg38, using identical alignment (bwa mem v0.7.17), filtering (MAPQ ≥ 30), SCALE normalization, SIP loop-calling parameters, and downstream counting with mariner (v1.5.0)**^31,41,42,45^**. Public ChIP-seq peak calls for transcription factors and histone marks in HEK293/HEK293T cells (GRCh38) were downloaded in bulk from the ENCODE portal (encodeproject.org). For each ENCODE experiment, the percentage of query peaks (retained/lost CTCF, retained/lost RAD21, or gained/lost ATAC-seq loop-anchor peaks) overlapping that experiment’s peak set was computed; experiments with multiple replicates targeting the same factor were averaged. When re-analyzing external RNA-seq or CUT&Tag datasets, we used the bagPipes workflows with the same tool versions and thresholds described above so that comparisons to our datasets were methodologically harmonized.

### Statistical Analysis

Statistical details for each experiment, including the statistical test used, exact value of n and what n represents, and the definition of center and dispersion/ precision measures (e.g., mean ± SEM), are reported in the corresponding figure legends.

## Declaration of generative AI and AI-assisted technologies

During the preparation of this work the author(s) used ChatGPT in order to identify weaknesses, correct grammar, and brainstorm titles. After using this tool/service, the author(s) reviewed and edited the content as needed and take(s) full responsibility for the content of the published article.

## Supplementary Figures

**Fig. S1.**
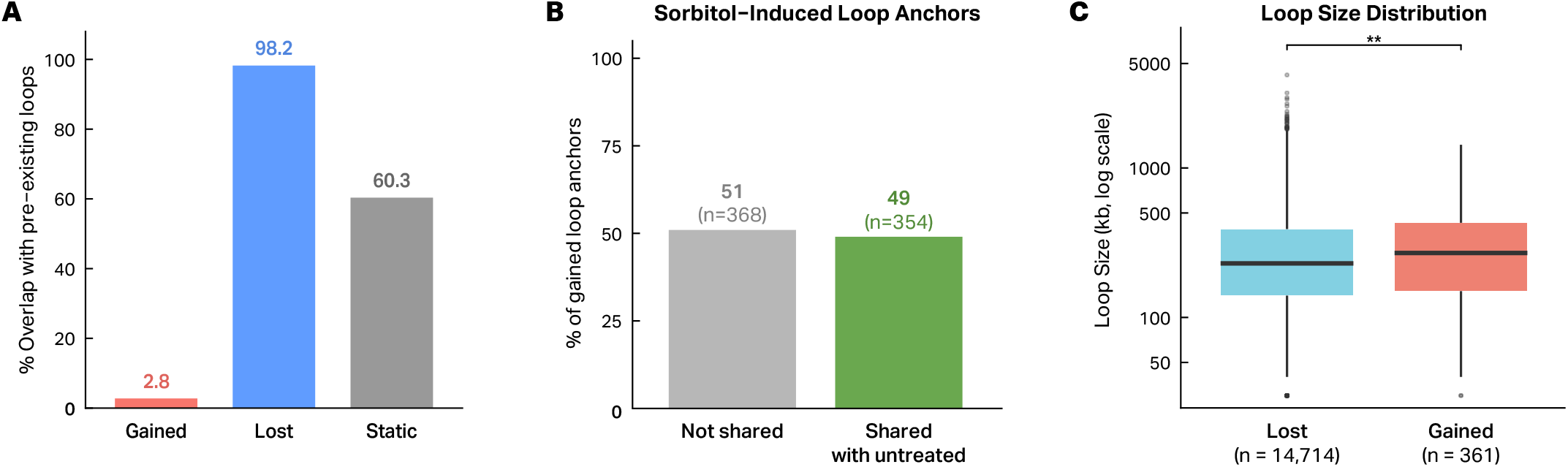
Hyperosmotic stress induces predominantly de novo looping that frequently reuses pre-existing anchor sites and favors long-range interactions. **A**, Percent overlap between differential loop categories and loop calls in untreated cells. Only 2.8% of sorbitol-induced (gained) loops overlap pre-existing loops. **B**, Proportion of sorbitol-induced loop anchors that coincide with loop anchor sites present in untreated cells. 49% of gained loop anchors (n = 354/722) overlap untreated anchor positions, whereas 51% are unique to sorbitol-treated cells. **C**, Size (kb) of lost and gained loops. Gained loops are significantly longer than lost loops (Wilcoxon rank-sum test; **p < 0.01),

**Fig. S2.**
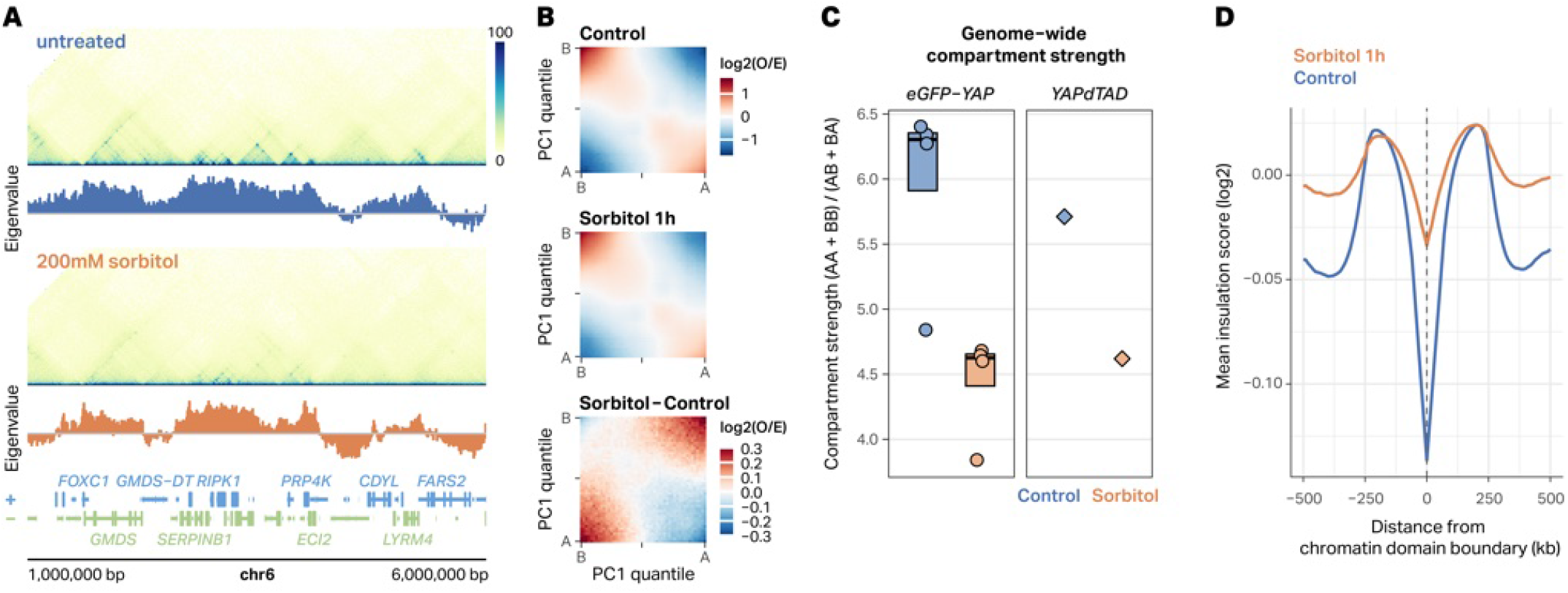
Hyperosmotic stress weakens A/B compartmentalization and chromatin domain insulation. **A**, Representative Hi-C contact maps (SCALE normalization, 10-kb resolution) at a 5-Mb region of chr6 (1–6 Mb, hg38) in untreated (top) and 200 mM sorbitol-treated (bottom) HEK293T eGFP-YAP1 cells, with corresponding PC1 eigenvector tracks shown below each map. Positive eigenvector values (blue) correspond to A compartment; negative values (orange) correspond to B compartment. **B**, Genome-wide saddle plots showing log2(observed/expected) contact frequency binned by PC1 quantile (50 bins) in control (top), sorbitol-treated (middle), and differential (sorbitol − control; bottom) conditions. Corners correspond to AA (active– active) and BB (inactive–inactive) interactions. **C**, Per-replicate compartment strength, quantified as (AA + BB) / (AB + BA) using the three corner bins of each saddle plot, in eGFP-YAP1 (n = 4 replicates per condition) and YAPdTAD cells (n = 1 per condition). Boxes show median and interquartile range; whiskers extend to 1.5× IQR. **D**, Mean insulation score (log2) centered on chromatin domain boundaries (±500 kb, 10-kb bins) in control (blue) and sorbitol-treated (orange) cells. The dashed vertical line marks the domain boundary position.

**Fig. S3.**
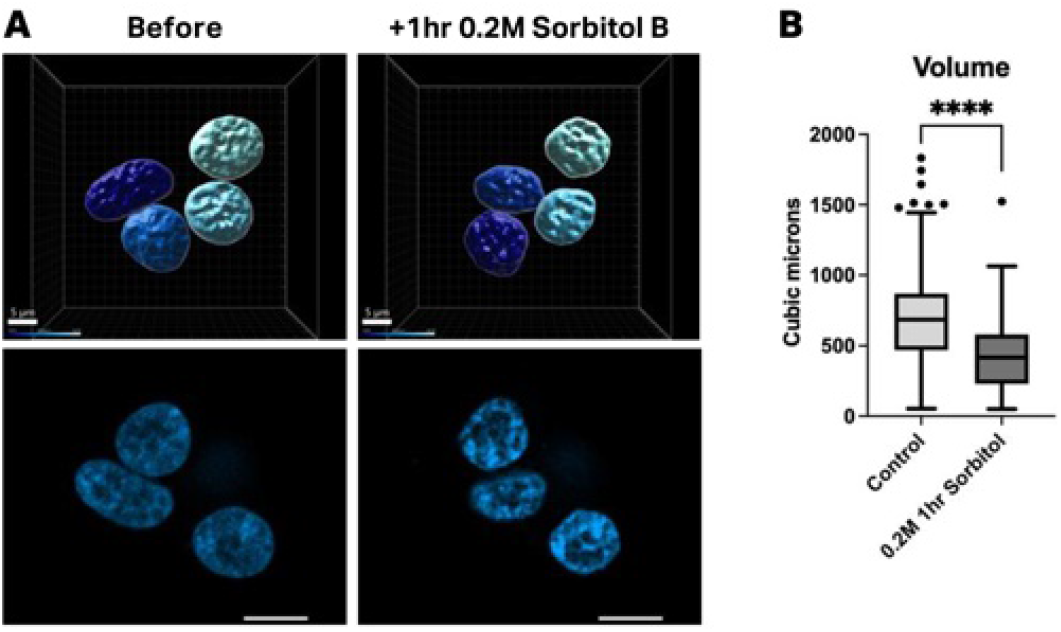
Hyperosmotic stress reduces nuclear volume. **A**, Representative 2D and 3D reconstructions of Hoechst-stained HEK293T nuclei before and after 1 h treatment with 0.2 M sorbitol. Scale bars, 10 μm. **B**, Quantification of nuclear volume measured from three-dimensional confocal image stacks. Each point represents an individual nucleus (Control, n = 416; Sorbitol, n = 436). Boxplots indicate median and interquartile range; whiskers extend to 1.5× IQR. Statistical significance was determined using a two-tailed Welch’s t-test (****P < 0.0001).

**Fig. S4.**
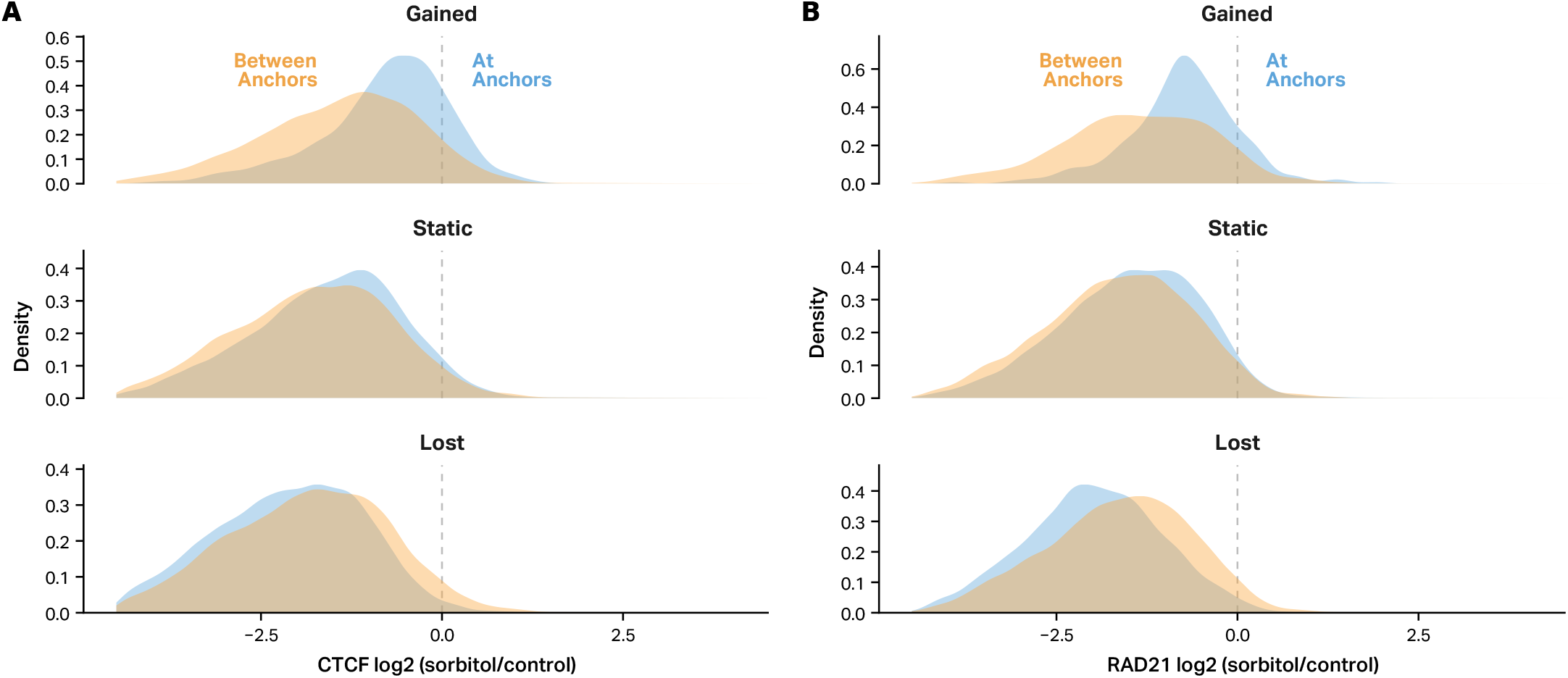
CTCF and cohesin binding is selectively retained at anchors of sorbitol-induced chromatin loops. **A**, Density distributions of CTCF log_**2**_fold–change values (sorbitol/control) at loop anchors (blue) versus the genomic regions between loop anchors (orange), stratified by loop class (static, lost, and gained). **B**, Density distributions of RAD21 log2fold-change values (sorbitol/control) at loop anchors (blue) versus the genomic regions between loop anchors (orange), stratified by loop class (static, lost, and gained).

**Fig. S5.**
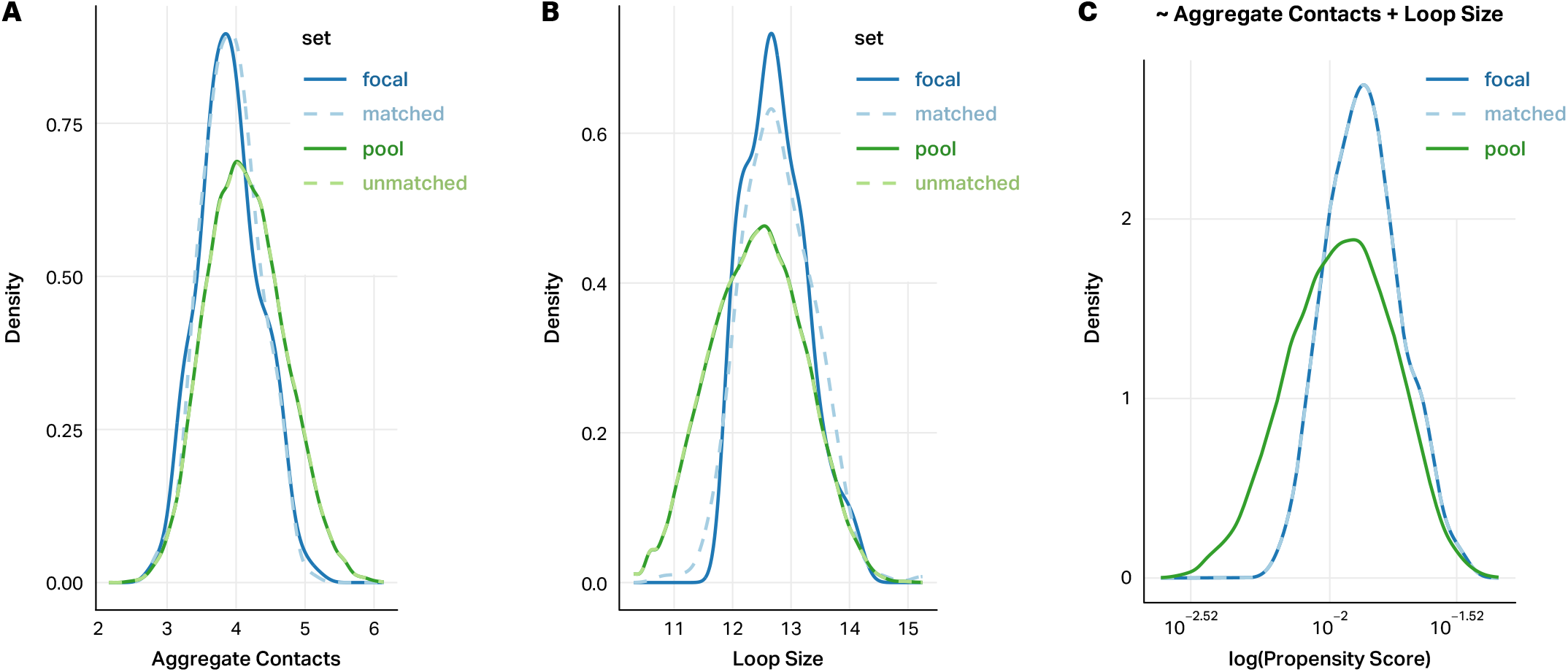
Covariate matching controls for loop size and interaction frequency in gained loop comparisons. **A**, Density distributions of aggregated Hi-C contact counts at loop anchors for focal gained loops (solid blue), the matched null set (dashed blue), and the full pool of candidate control loops (solid/dashed green). **B**, Density distributions of loop size (log-transformed genomic span) for the same loop sets. **C**, Propensity score distributions for focal, pool, and matched loops (aggregated contacts and loop size).

**Fig. S6.**
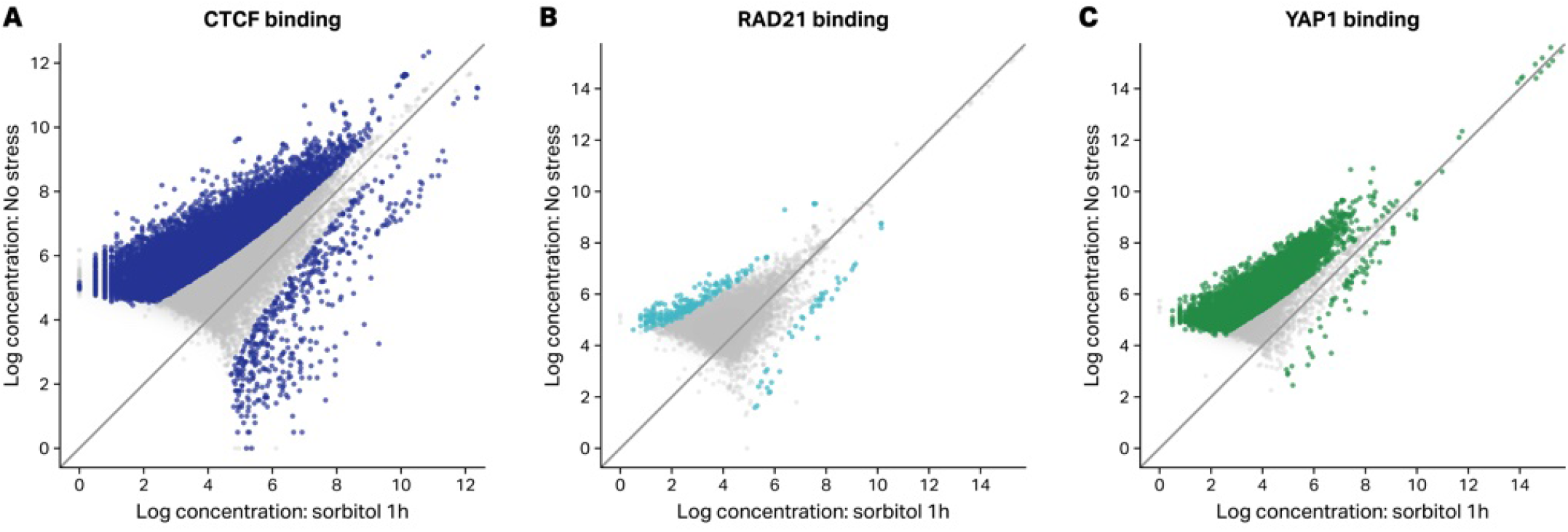
Genome-wide CUT&Tag signal shows predominantly reduced occupancy for CTCF, RAD21, and YAP1 after hyperosmotic stress. Scatter plots of log2-transformed mean CUT&Tag counts at called peaks for (**A**) CTCF, (**B**) RAD21, and (**C**) YAP1, comparing no-stress to 1h sorbitol-stress conditions. Each point represents a single peak; colored points are statistically significant (DESeq2, padj < 0.05 for CTCF and YAP1; padj < 0.1 for RAD21), and gray points are non-significant.

**Fig. S7.**
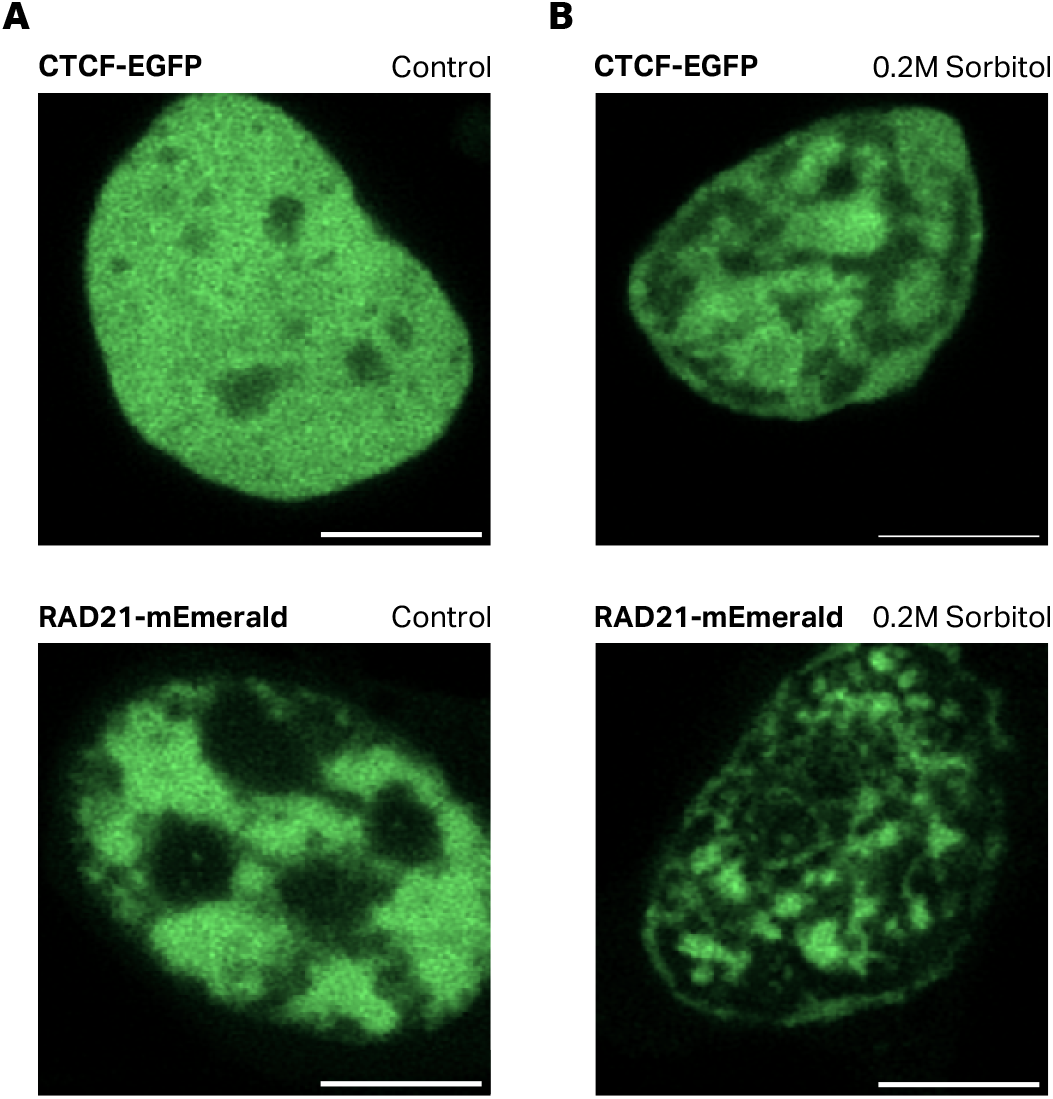
CTCF and RAD21 redistribute following hyperosmotic stress. **A**, Representative fluorescence images of untreated CTCF-eGFP HEK293 cells (top) and untreated RAD21-mEmerald HEK293 cells (bottom). **B**, Representative fluorescence images of CTCF-eGFP HEK293 cells (top) and RAD21-mEmerald HEK293 cells (bottom) treated with 0.2 M sorbitol. Scale bars, 5 μm.

**Fig. S8.**
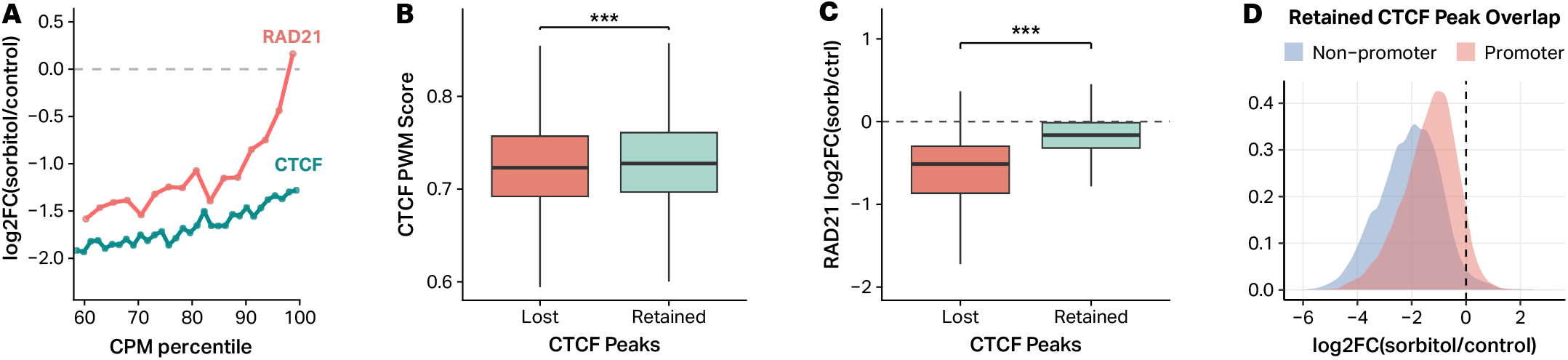
CTCF retention under hyperosmotic stress is associated with higher baseline occupancy, stronger CTCF motifs, co-retention of RAD21, and promoter overlap. **A**, Median log2 fold change (sorbitol/control) in CUT&Tag signal for CTCF (teal) and RAD21 (red) as a function of binding occupancy in untreated cells, measured as counts per million (CPM) percentile bin (bins 60–100 shown). **B**, CTCF position weight matrix (PWM) scores at retained versus lost CTCF peaks, scored within a 50-bp window centered on the CTCF summit. Retained peaks have significantly higher PWM scores than lost peaks (Wilcoxon rank-sum test, ***p < 0.001). Boxes show median and interquartile range; whiskers extend to 1.5× IQR. **C**, RAD21 log2 fold change (sorbitol/control) at CTCF peak sites, stratified by CTCF retention status (Wilcoxon rank-sum test, ***p < 0.001). Boxes show median and interquartile range; whiskers extend to 1.5× IQR. **D**, Density distributions of CTCF CUT&Tag log2 fold change (sorbitol/control) at promoter-overlapping (red) versus non-promoter (blue) CTCF peaks.

**Fig. S9.**
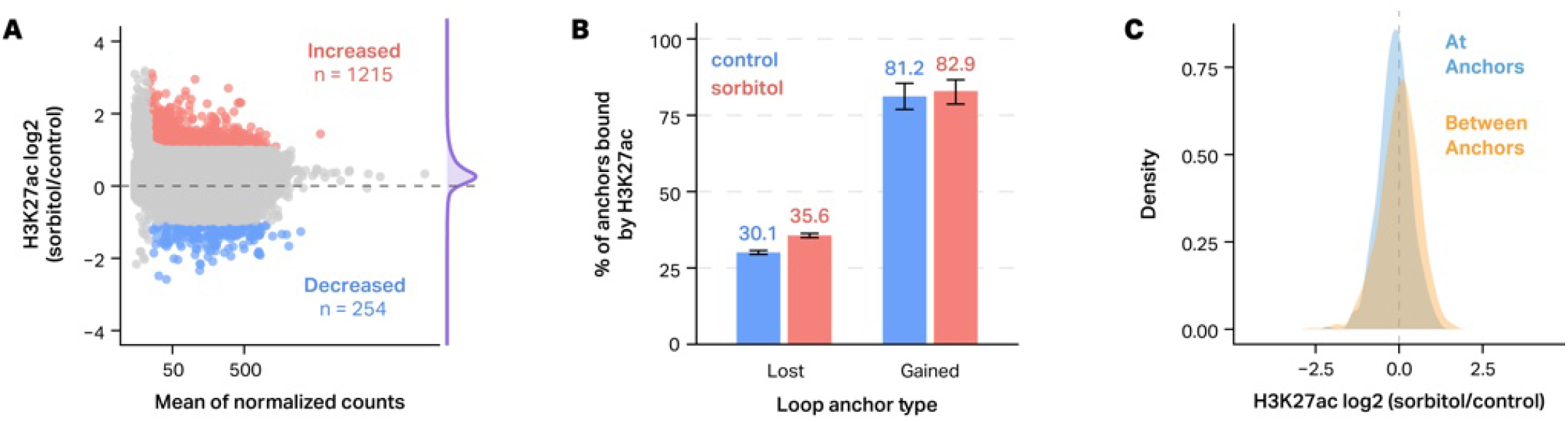
Hyperosmotic stress differentially modulates H3K27ac binding at loop anchors. **A**, MA plots showing differential CUT&Tag signal for H3K27ac in sorbitol-treated versus control HEK293T cells. The x-axis denotes the mean of normalized counts and the y-axis shows log**2** occupancy (sorbitol/control); peaks with significantly increased or decreased signal (DESeq2, padj < 0.05) are highlighted in red and blue, respectively, with the number of sites indicated. **B**, Percentage of lost and gained loop anchors that are bound by H3K27ac in control (blue) and sorbitol-treated (red) cells. **C**, Density plots of H3K27ac occupancy changes at gained loop anchors, comparing peaks at loop anchors (blue) versus the regions between anchors (orange).

**Fig. S10.**
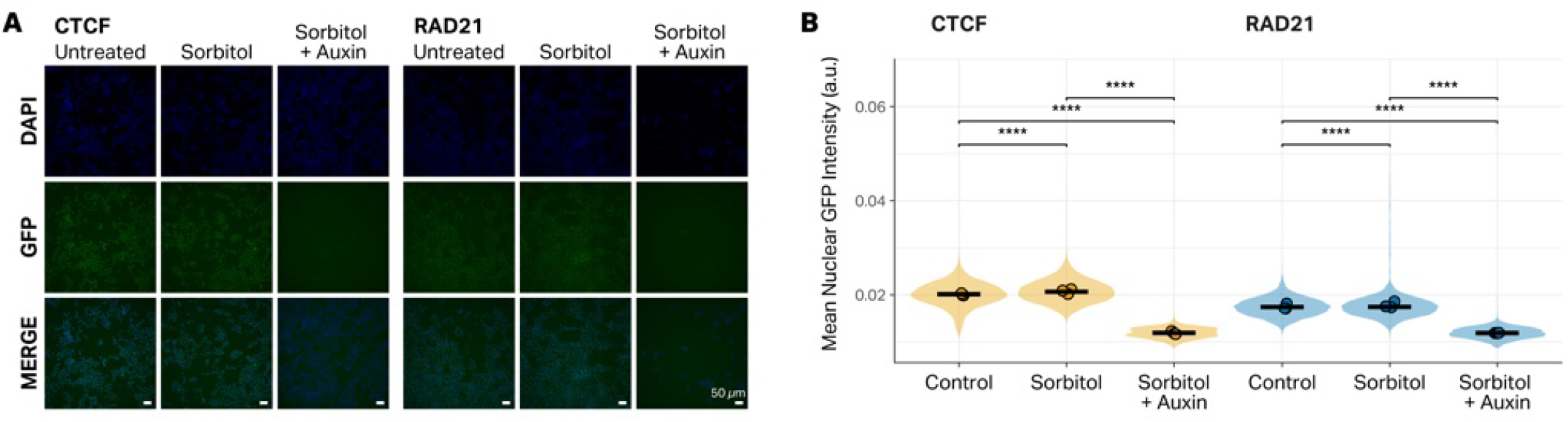
Auxin-inducible degradation efficiently depletes CTCF and RAD21 in HCT116 cells. **A**, Representative fluorescence microscopy images of HCT116-CTCF-mAID2 (left) and HCT116-RAD21-mAID2 (right) cells under three conditions: untreated, 200 mM sorbitol (1 h), and sorbitol + 1 µM 5-Ph-IAA auxin (1 h). Rows show DAPI (blue), GFP (green), and merged channels. Scale bars, 50 μm. **B**, Quantification of mean nuclear GFP intensity (arbitrary units) from CellProfiler segmentation, shown as violin plots with per-field replicate means overlaid (dots) and median crossbars. Welch’s t-tests are shown for all three condition comparisons within each protein (****p < 0.0001).

**Fig. S11.**
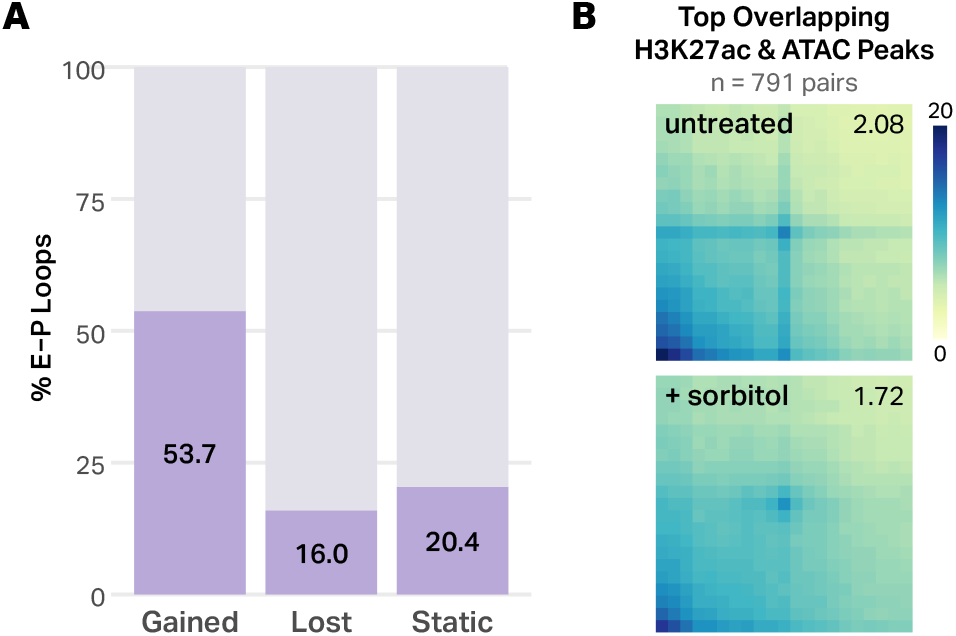
Sorbitol-induced loops are enriched for enhancer-promoter interactions while global contacts between H3K27ac sites slightly decrease in response to sorbitol. **A**, Proportion of gained, lost, and static loops classified as enhancer-promoter (E-P) interactions. Loops were defined as E-P if one anchor overlapped an annotated promoter (derived from TxDb hg38) and the distal anchor overlapped an H3K27ac peak not overlapping a promoter. Gained loops show the highest E–P fraction (53.7%) compared to lost (16.0%) and static (20.4%) loops. **B**, Aggregate Peak Analysis (APA) of Hi-C contact frequency between pairs of cis-regulatory elements (CRE). CREs were defined as loci with overlapping H3K27ac and ATAC peaks. Interaction frequency was quantified between the 1,000 CRE with the highest occupancy of H3K27ac, filtered for pairs of CRE separated by 25 kb to 1 Mb (n = 791 pairs). Matrices show aggregate observed contact frequency (10-kb resolution) in untreated (top) and sorbitol-treated (bottom) HEK293T eGFP-YAP1 cells. APA scores (foreground/background enrichment) are indicated in the upper-right corner of each matrix.

**Fig. S12.**
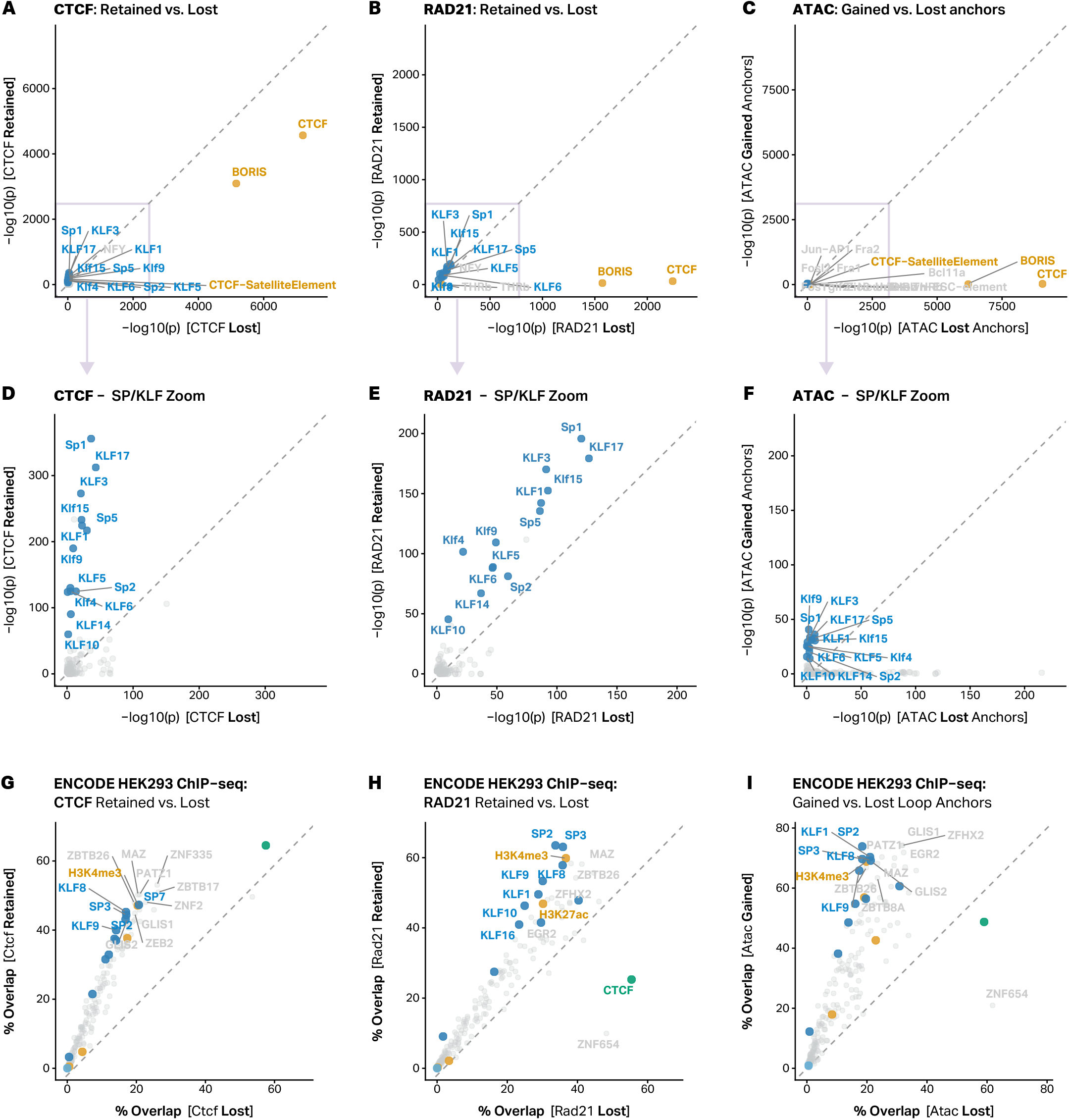
Binding sites for most TFs are enriched at retained CTCF and RAD21 sites and at gained loop anchors. **A–C**, HOMER motif enrichment QQ scatter plots comparing the significance (−log10 p-value) of known motifs between retained and lost CTCF peaks (**A**), retained and lost RAD21 peaks (**B**) and gained versus lost loop anchors overlapped with ATAC-seq data (**C**). Each point represents one motif; CTCF/BORIS family motifs (teal) and KLF/SP family motifs (gold) are highlighted. The dashed diagonal indicates equal enrichment. **D**–**F**, SP/KLF zoom-in of panels A–C, respectively, with CTCF/BORIS motifs excluded to rescale axes. **G**–**I**, ENCODE HEK293/HEK293T ChIP-seq overlap scatter plots for the same three comparisons (CTCF retained vs. lost, RAD21 retained vs. lost, and ATAC gained vs. lost anchors, respectively). Each point represents one factor or histone mark; the percentage of peaks from each comparison category overlapping that factor’s peaks is shown on each axis.

**Fig. S13.**
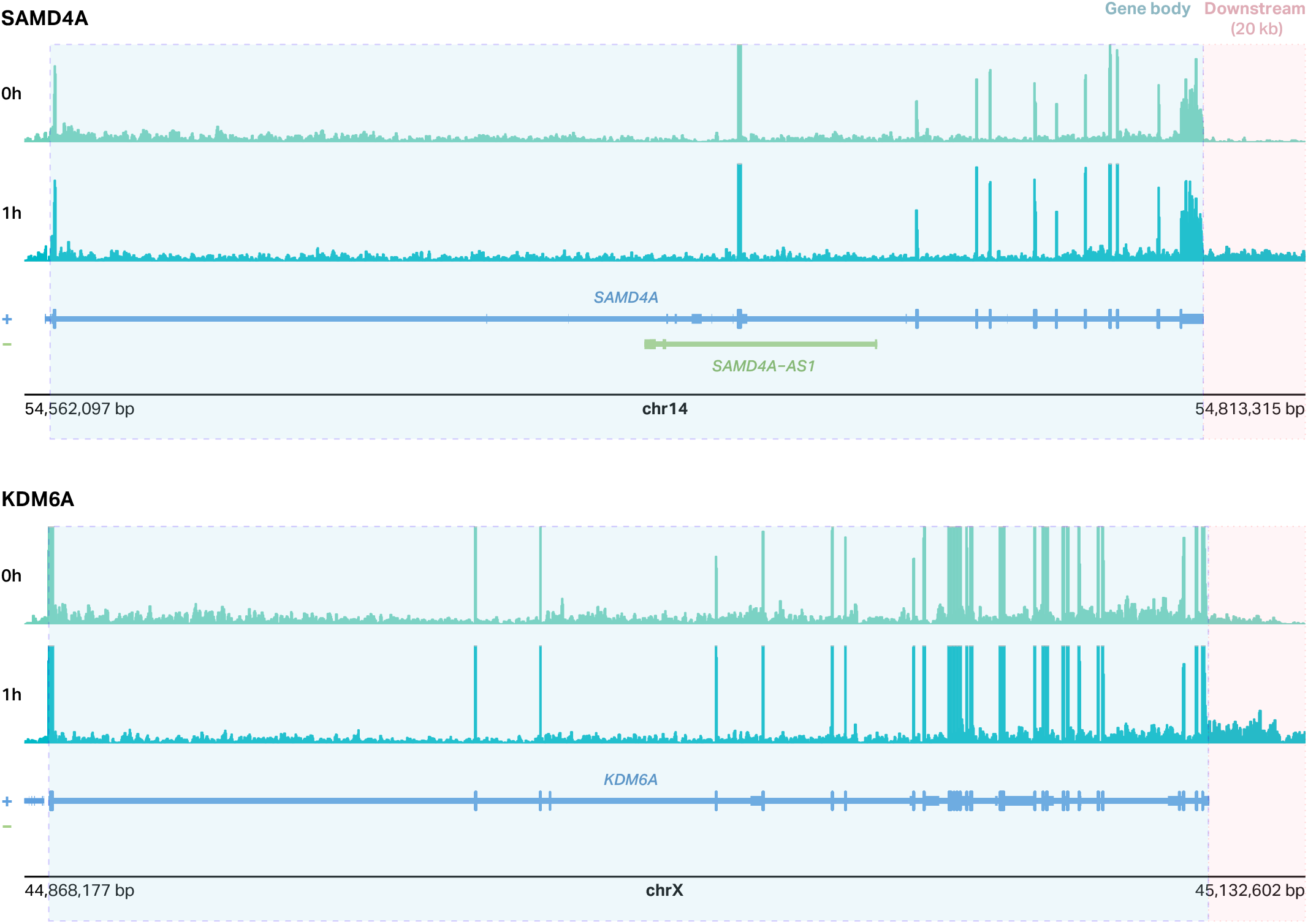
Hyperosmotic stress induces downstream-of-gene (DoG) transcription at select loci. Genome browser views of RNA-seq signal at the SAMD4A (top) and KDM6A (bottom) loci in HEK293T cells before (0 h) and after 1 h of 200 mM sorbitol treatment. The annotated gene body is highlighted in blue and the 20-kb downstream region used for DoG detection is shaded in pink.

